# Evidence for impaired hippocampal circuitry in schizophrenia and its link to memory dysfunction

**DOI:** 10.1101/2023.11.05.565219

**Authors:** Asieh Zadbood, Yingying Tang, Wenjun Su, Hao Hu, Gillian Capichioni, Shuwen Yang, Junjie Wang, Camille Gasser, Oded Bein, Li Hui, Qiufang Jia, Tianhong Zhang, Yawen Hong, Jijun Wang, Donald Goff, Lila Davachi

**Affiliations:** Department of Psychology, Columbia University, New York; Shanghai Mental Health Center, Shanghai Jiao Tong University School of Medicine, Shanghai, China; Department of Psychiatry, New York University Langone Medical Center, New York; Department of Developmental and Behavioural Pediatric & Child Primary Care, Xinhua Hospital, Shanghai Jiao Tong University School of Medicine, Shanghai, China; Suzhou Guangji Hospital, The Affiliated Guangji Hospital of Soochow University, Suzhou, China; Princeton Neuroscience Institute, Princeton University, New Jersey; Nathan Kline Institute for Psychiatric Research, Orangeburg, New York

## Abstract

Pattern separation and pattern completion are opposing yet complementary components of mnemonic processing that heavily rely on the hippocampus. It has been shown that processing within the dentate gyrus (DG) subfield promotes pattern separation while operations within the CA3 subfield are important for pattern completion. Schizophrenia has been associated with anatomical and functional hippocampal abnormalities, including within the DG and CA3. We hypothesized that an impairment in hippocampal circuitry in individuals with first-episode schizophrenia leads to deficits in pattern separation (mnemonic discrimination) and pattern completion (recognition memory), that these deficits contribute to delusions, and that antipsychotic treatment improves circuit functioning. We measured behavioral and neural responses during the identification of new, repeated, and similar stimuli using high-resolution fMRI in 45 medication-free or minimally-treated patients with first-episode schizophrenia and 49 matched controls. We found recognition memory and pattern separation deficits in patients and a negative association between memory performance and the severity of delusions. Neural analyses revealed deficits in both univariate BOLD responses and multivariate patterns in the hippocampus during mnemonic discrimination in patients compared to controls. Importantly, by investigating the association between trial-level neural activity and behavior before and after treatment, we found that antipsychotics normalized DG activity during pattern separation and CA3 activity during pattern completion. Lastly, trial-level cortical responses during mnemonic discrimination predicted performance in patients at baseline, suggesting a compensatory role. This study provides new insight into the impact of schizophrenia and antipsychotic treatment on memory systems and uncovers systems-level contributions to pattern separation and pattern completion.

## Introduction

An intact memory system is essential for cognition. Memories are not just references to the past but building blocks of the current thoughts and beliefs that shape expectations and the experience of reality. Thus, deficits or alterations in memory function may not only lead to forgetting but also to disorders of thought and distortions of reality. Schizophrenia is characterized by psychosis, which is an impairment in the interpretation of reality. Accumulating evidence supports a potential link between schizophrenia and memory system abnormalities ^1–4^. Individuals with schizophrenia exhibit deficits in relational memory ^5,6^ and item recollection ^7,8^, may form fixed false memories ^9^ (associated with delusions ^10^), and are impaired in mnemonic discrimination of highly similar events ^11,12^. Accordingly, several lines of evidence point to the hippocampus as a critical region affected by the disorder ^3,13^, strengthening the hypothesis that hippocampal dysfunction may play a role in the pathophysiology of schizophrenia^1,2,3,14^. There has been consistent evidence for hippocampal volume reduction ^13,15,16^ prominent in the early stages of the illness ^17^, which is more pronounced than reductions in other brain structures ^18^. Several studies have demonstrated impaired connectivity between the hippocampus and other cortical and subcortical areas ^19,20^, and abnormal activation of the hippocampus during memory tasks ^6,21–23^. Given the complex nature of hippocampal circuitry, however, it is unclear what role hippocampal circuits may play in memory impairment and reality distortion in schizophrenia.

Prior work on hippocampal circuitry in schizophrenia has reported impairments in hippocampal subfields, in particular the dentate gyrus (DG) and cornu ammonis 3 (CA3) ^24–26^. Sparse coding in the DG is hypothesized to be an important mechanism for orthogonalizing neural representations of highly similar stimuli, known as *pattern separation* ^27,28^. This process is believed to allow for successful mnemonic discrimination -- the ability to distinguish new from formerly seen similar items. Moreover, DG is a site of neurogenesis that plays a role in pattern separation^29^ and is highly sensitive to environmental stressors early in development ^30^. These environmental stressors have also been identified as risk factors for schizophrenia, including hypoxia^31^, inflammation^32^, and associated oxidative stress ^33^. Accordingly, reduced neurogenesis has been reported in schizophrenia ^34^. The DG receives input from the entorhinal cortex and sends outputs to CA3 via mossy fibers; CA3 also receives direct input from the entorhinal cortex. It is thought that the auto-associative neural architecture of CA3 plays a significant role in *pattern completion* by strengthening mnemonic associations and recovering associations from partial input, which is essential for memory retrieval ^35,36^. Reduction in mossy fiber synapses has been reported in schizophrenia, ^37,38^ potentially leading to a loss of inhibitory input to CA3 ^1^. This has led to the hypothesis that hypoactivity in DG and hyperactivity in CA3 may be a core cause of memory dysfunction in schizophrenia ^1^.

Dysfunction in the DG/CA3 circuitry may contribute to the generation and persistence of delusions through impaired discrimination between new and past experiences (impaired pattern separation due to DG dysfunction) and the strengthening of these aberrant associations (CA3 dysfunction). Some theoretical accounts of delusional ideation suggest a link between delusions and impairments in the associative memory system^1,39^. However, to our knowledge, no empirical study to date has directly examined the function of the DG/CA3 circuit during mnemonic judgments that differentially tax pattern separation and completion in individuals with schizophrenia. We hypothesized that impairments specifically in the DG/CA3 circuitry might disrupt the balance between pattern separation and pattern completion and lead to decreased mnemonic discrimination (pattern separation) and impaired recognition memory (pattern completion) in individuals with schizophrenia. We predict that these memory impairments will be higher in individuals with more severe delusions. We reasoned that if antipsychotics improve hippocampal integrity ^40^, we might observe a regularization in DG/CA3 circuit function in patients after treatment (Figure 1-A).

**Figure 1:**
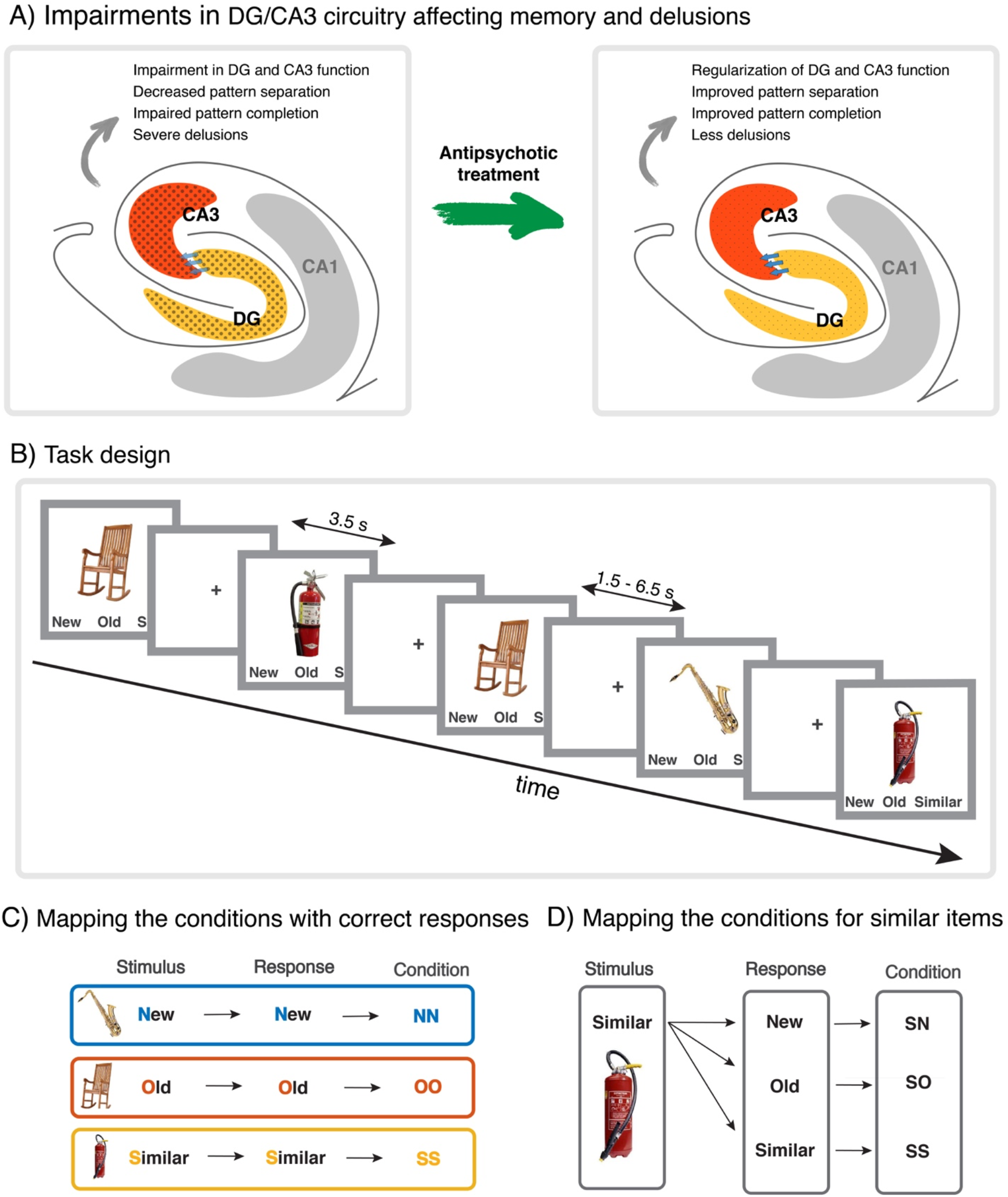
A) The hypothesized role of the DG and CA3 impairments in memory behavior and delusion severity in individuals with schizophrenia and the effect of antipsychotic treatment on regularizing the circuit function. B) Task design. C) Illustrates the mapping of trial types and responses during correct trials. NN denotes a “new” response to new trials. OO denotes “old” response to old trials. SS denotes a “similar” response to similar trials. The same color codes will be used throughout the paper to refer to the results for the new (blue), old (red) and similar (orange) stimuli. D) shows the mapping of trial type and responses during similar trials. SN denotes a “new” response to similar trials (incorrect response). SO denotes an “old” response to similar trials (incorrect response). SS shows a “similar” response to similar trials (correct response).

We tested these hypotheses using high-resolution fMRI while participants performed a widely used Mnemonic Similarity Task (MST) ^41^. The MST allows measurement of pattern separation and pattern completion because it requires participants to recognize when an item is old and discriminate that from when an item is very similar, but not identical, to one already presented. We implemented a continuous recognition version of the task in which participants identified each item as “new”, “old”, or “similar” (Figure 1-B/C). A few empirical studies have used the MST to measure mnemonic discrimination behavior (no neural data) in chronic and first-episode schizophrenia ^11,12,42^. In these studies, pattern separation performance was decreased in individuals with schizophrenia, consistent with DG pathology. However, recognition memory performance was inconsistent across these studies. Additionally, behavioral differences between patients and controls have not been consistently linked with clinical measures of illness. We studied newly diagnosed patients with delusions and a diagnosis of schizophrenia with little or no prior antipsychotic treatment, as prior research suggested disease progression and antipsychotic treatment can affect both the function and structure of the hippocampus ^40,43,44^. We measured both univariate and multivariate BOLD signals in the DG, CA3, and CA1 subfields while participants were making mnemonic judgments and asked, on a trial-by-trial basis, whether the neural activity predicted mnemonic judgment success before and after treatment with antipsychotics. Taken together, to the best of our knowledge, we provide the first evidence for the functional impairment of hippocampal circuitry in schizophrenia and its link to memory function, delusions, and antipsychotic efficacy.

## Results

### Behavior during the MST task

To evaluate recognition memory behavior, we calculated the proportion of “old” responses to old stimuli (OO) and controlled for response biases, similar to previous work ^11,12,42^. Specifically, to correct for any general bias to report an item as “old”, we subtracted NO (responding “old” to new stimuli), from OO. Using this *recognition memory index*, we found that recognition memory was reduced in patients compared to controls (t(92) = −3.66, 95%CI [−0.24 −0.07], p = 0.0004, Figure 2-A). Because the MST task places strong demands on the retrieval of specific details to discriminate an old from a similar trial, we computed an additional metric to look at overall recognition memory for old stimuli. Specifically, we reasoned that both “old” and “similar” responses to an old stimulus convey that participants recognize the item as being familiar. Thus, we calculated an overall recognition score by considering both “old” or “similar” responses to an old stimulus while controlling for general biases to report “old” or “similar” (OO + OS – NO – NS). Using this metric, we observed that the overall recognition of old stimuli was reduced in patients compared to healthy controls (t(92) = 3.73, 95%CI [−0.20 −0.06], p = 0.0003, Figure 2-B).

**Figure 2:**
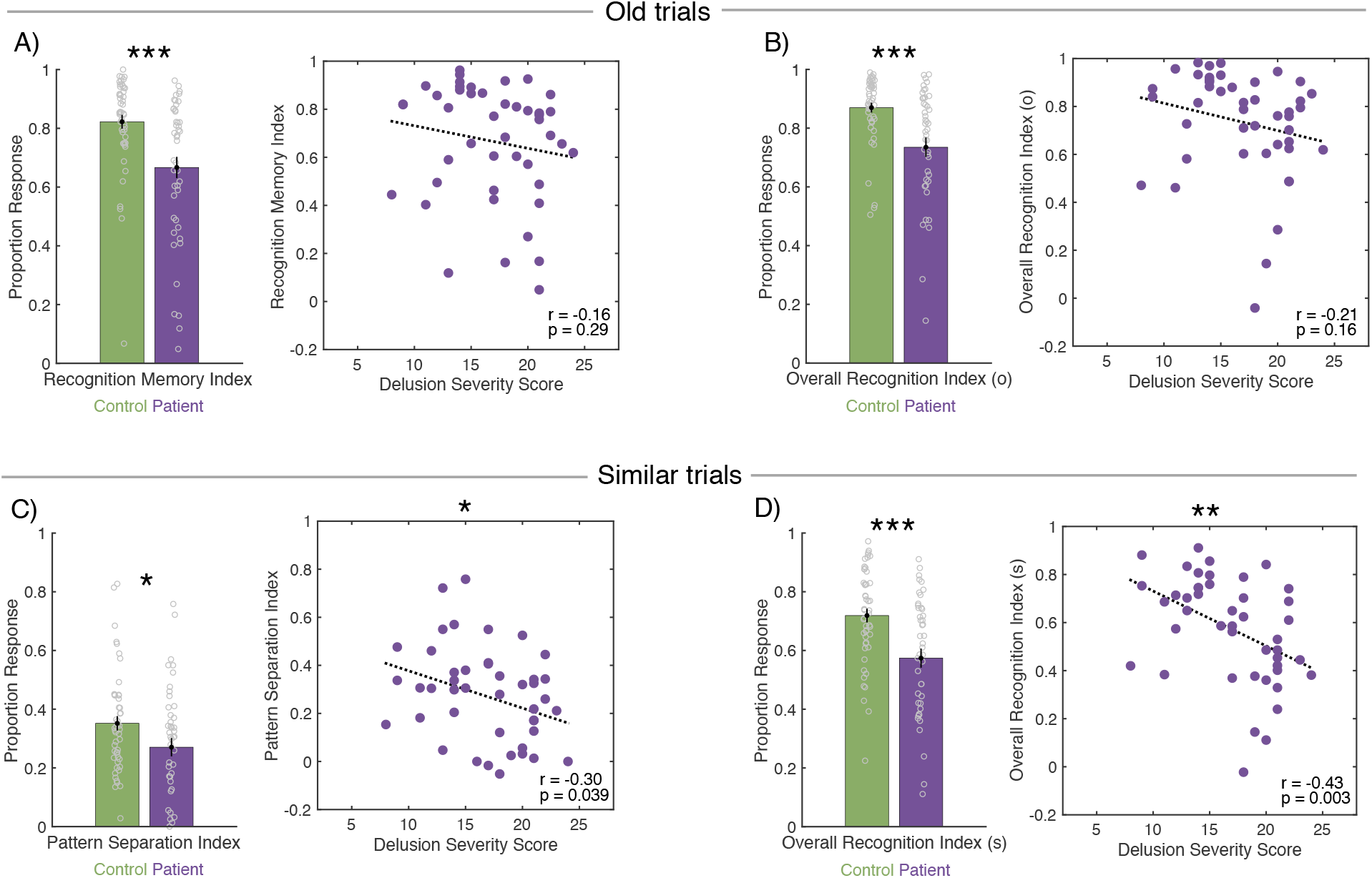
Behavioral performance in Session 1. **A)** Recognition memory index (OO – NO) in patients and controls and the relationship between recognition memory index and delusions severity score in patients. Overall recognition index for old trials (OO + OS – NO – NS) in patients and controls and the relationship between overall recognition index for old trials and delusions severity score in patients. **C)** Pattern separation index (SS - NS) in patients and controls and the relationship between pattern separation index and delusion severity in patients. **D)** Overall recognition index for similar trials (SS + SO – NO – NS) in patients and controls and the relationship between overall recognition index for similar trials and delusions severity score in patients.

Now we turn to similar trials to examine the mnemonic discrimination behavior. In keeping with prior reports using the MST task ^11,12,42^, we calculated a *pattern separation index*; which corrects for response biases by subtracting the proportion of times participants use the “similar” response to new trials (NS) from their correct rate of response to similar trials (SS). Using this index, we found a significantly smaller pattern separation index in patients compared to controls during Session 1 (t(92) = −2.07, 95%CI [−0.15 −0.003], p = 0.04, Figure 2-C). Then we computed an overall recognition index (as above) but during responses to similar trials. We counted the proportion of both “old” and “similar” responses to similar stimuli and subtracted the general bias to respond “old” or “similar” (SS + SO – NS - NO). Patients also showed a significant reduction in their overall recognition of similar stimuli compared to healthy controls (t(92) = −3.66, 95%CI [−0.22 - 0.06 0.1], p = 0.0004, Figure 2-D).

Further examination of the general bias to respond “old” or “similar” (NO + NS), revealed that this bias is especially high in patients. We report this index as a potential indicator of *excessive pattern completion*, which was larger in patients in Session 1 (t(92) = 2.07, p = 0.04, 95%CI [0.002 0.11) and significantly dropped in Session 2 after treatment (t(28) = 2.91, p = 0.006, 95% CI [0.02 0.16]).

#### The relationship between behavioral mnemonic discrimination and delusion severity

We observed an association between the pattern separation index and delusion severity, assessed by PSYRAT, where patients with higher index scores had less severe delusions in Session 1 (r = - 0.30, p = 0.039, Figure 2-C). Importantly, there was a significant correlation between delusion severity and the deficit in overall recognition of similar trials. Specifically, patients with more severe delusion symptoms -- measured by PSYRAT delusion score – exhibited a lower overall recognition index (r = −0.43, p = 0.003, Figure 2-D). There was no significant relationship between the delusion severity score and recognition memory index (r = −0.16, p = 0.29, Figure 2-A) or overall recognition of old stimuli (r = −0.21, p = 0.16, Figure 2-B).

Comparing the performance between Session 1 and 2, we did not observe significant differences in the recognition memory indices or the pattern separation index in either patients or controls (see supplementary results for a full report of Session 2). However, after antipsychotic treatment, the delusion severity scores significantly dropped (t(28) = 14.62, 95%CI [12.84 17.02], p = 1.2e- 14; Session 2 score equal to zero in 22 out of 29 patients), and thus, there was not enough variability in data to assess the relationship between delusion severity and behavior in Session2.

### Hippocampal subfield BOLD activation during the MST task

#### Univariate BOLD responses in hippocampal subfield regions of interest (ROIs)

The majority of prior work using the MST utilized it in the context of an incidental encoding paradigm where participants were not required to discriminate presented items ^45–47^. Specifically, Bakker et al ^45^ used the incidental encoding MST, in which participants viewed new, old, and similar items and made indoor/outdoor judgments. They showed that in a *combined* DG/CA3 subregion, but not in CA1 or subiculum, the BOLD activity in response to similar items was higher than in old trials and was comparable to encoding new items (which may suggest lack of repetition suppression during similar trials). In an earlier study^48^ in which participants made memory judgments for new, old, and similar stimuli, however, BOLD activation during correct identification of old and similar items did not differ in the combined DG/CA3 ROI. They showed that the BOLD response during SS was numerically lower than during OO in DG/CA3 and significantly lower than OO response in CA1. We, too, implemented a version with explicit memory demands to obtain mnemonic behavior during BOLD acquisition, in light of the reports on behavioral pattern separation deficits in individuals with schizophrenia^11,12,42^ (which we also replicated, see behavioral results). Using this task allowed us to isolate our analyses to correct similar and old responses, where we were more certain successful recognition and/or mnemonic discrimination had occurred. In addition, we were able to use trial-level memory responses (either correct or incorrect) to model the association between the BOLD activation and behavioral performance – a novel approach to analyzing the MST data (see trial-level results). Importantly, unlike most prior work in humans^45–48^, we investigated pattern separation and pattern completion in separate DG and CA3 ROIs.

We first computed the BOLD percent signal change in DG, CA3, and CA1 subregions of the hippocampus during the correct identification of similar and old items (SS and OO)(for details see Methods). There was a significant main effect of condition (DG: F(1,81) = 7.95, p = 0.006, CA3: F(1,81) = 8.32, p = 0.005, Figure 3-A, B) and a significant interaction between group and condition (DG (F(1,81) = 6.08, p = 0.01, CA3 F(1,81) = 8.32, p = 0.005,). In the control group, pairwise comparisons revealed significantly lower BOLD activation during SS responses compared to OO, in both DG (p = 0.001, Cohen’s d = 0.64) and CA3 (p = 0.0007, Cohen’s d = 0.70). In patients, however, there were no significant differences between BOLD activation during SS and OO in DG or CA3. Direct comparisons between patients and control during SS trials revealed significantly lower percent signal change in controls in both DG (p = 0.018, Cohen’s d = 0.57) and CA3 (p = 0.008, Cohen’s d = 0.59). No significant difference was observed in BOLD activation during OO trials between the two groups. In CA1, the main effect of condition was not significant but a significant group*condition interaction was observed (F(1,81) = 5.09, p = 0.014, Figure S4-A). In a pairwise comparison, BOLD activation during SS was significantly lower than OO in controls (p = 0.018, Cohen’s d = 0.48), but not patients (p = 0.83). Direct comparison of the neural response during SS in patients and controls revealed a significantly lower response in controls (p = 0.011, Cohen’s d = 0.56). The neural response during correct identification of new trials (NN) is presented for comparison in the figures as we expected a higher response (novelty response) in this condition compared to SS and OO (Figures 3 and S4). However, the statistical analysis only included the main conditions of interest (SS and OO).

**Figure 3:**
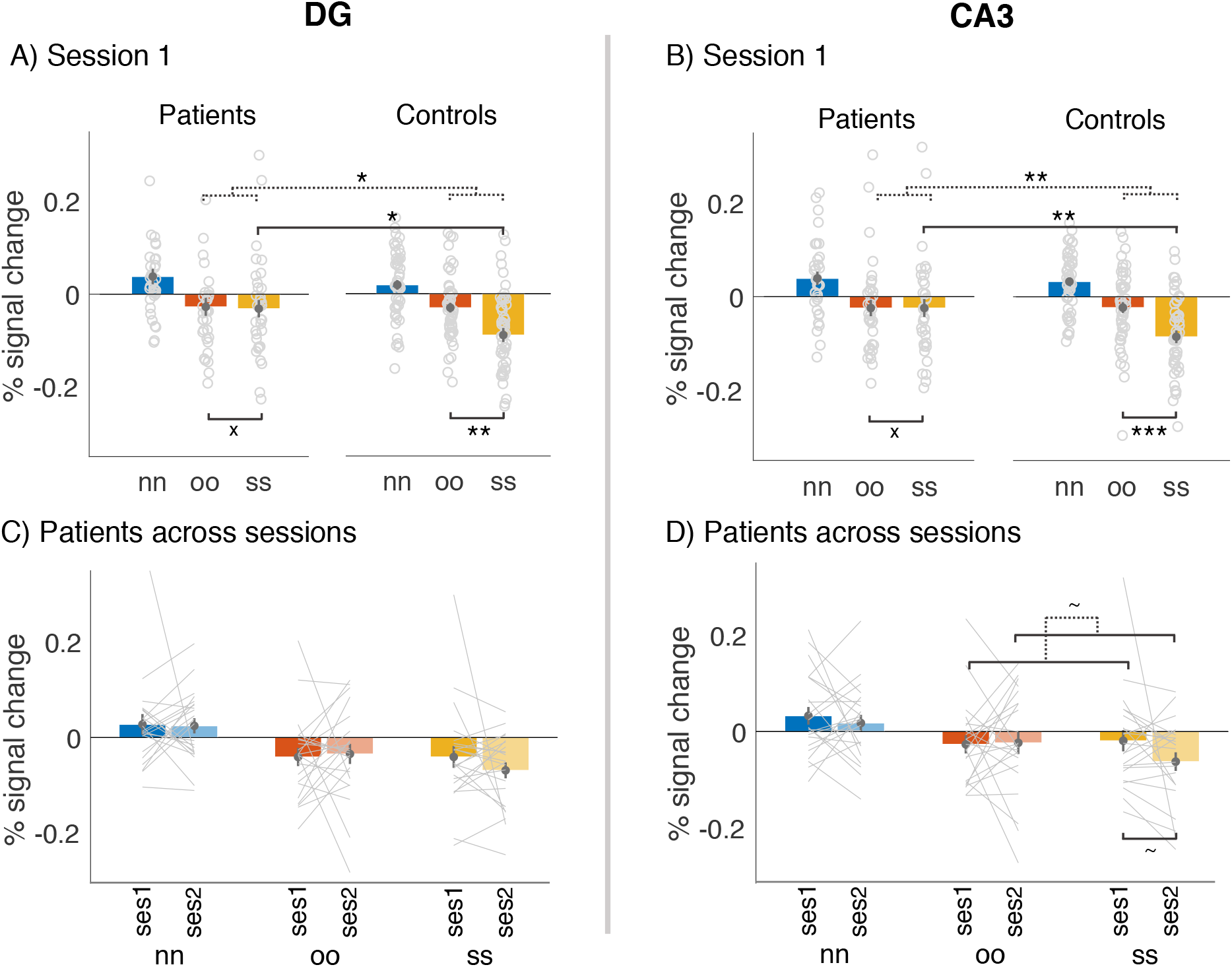
Percent signal change during correct responses to new, old, and similar conditions (NN, OO, SS). Dotted lines depict the interaction between the two linked comparisons. **A)** Neural response in DG in Session 1 across the two groups **B)** Neural response in CA3 in Session 1 across the two groups **C)** Neural response in DG across the two sessions in patients **D)** Neural response in CA3 across the two sessions in patients. Statistical analyses were performed on conditions of interest (SS and OO). NN is shown as a control condition.

The pattern of BOLD response across conditions remained the same in Session 2 in both patients and controls. We directly compared BOLD activity between Session 1 and Session 2 in the subset of participants who completed the follow-up session (see Methods for participant numbers in each analysis). We asked whether the BOLD response changed from Session 1 to Session 2 in either group and if the pattern of response in patients after medication treatment became more similar to what we found in controls (BOLD response to SS < OO). In patients, there was no main effect of condition or session and there was no significant difference between BOLD activity during SS and OO conditions after treatment, similar to what we observed in Session 1 (Figure 3- C, D). However, there was a trend toward a condition * session interaction in CA3 in patients (F(1,20) = 2.4, p = 0.13), driven by more negative BOLD response during SS condition in Session 2 compared to Session 1 (paired comparison: p = 0.14), suggesting that the response pattern in patients became more similar to controls (BOLD response to SS < OO). In controls, DG, CA3, and CA1 showed a significant main effect of condition (DG: F(1,45) = 62.1, p = 5.06e-10, CA3: F(1,45) = 43, p = 4.63e−08, CA1: F(1,45) = 38.7, p = 1.46e−07), driven by more negative responses in SS compared to OO condition in Sessions 2 (paired t test - DG: p = 0.0002, CA3 = 0.0004, CA1 = 0.009) similar to Session 1 (paired t test - DG: p = 0.0008, CA3 = 0.001, CA1 = 0.0004) (Figure S5). Lastly, no significant difference was found in the BOLD response to SS between two sessions in controls.

#### Multivariate neural pattern analyses in hippocampal subfield ROIs

More recently, multivariate pattern analyses have been used to study pattern separation, where an index of ‘separation’ would manifest as more differentiated neural patterns for the similar condition compared to the old condition ^49–51^. After showing a robust difference in the averaged neural response to SS and OO conditions between patients and controls, we measured more fine-grained spatial patterns of BOLD activation in these ROIs during different memory conditions. To this end, we extracted beta estimates for each trial in each voxel and created vectors of beta estimates (neural patterns) for each trial in each ROI (LSS approach ^52^, see Methods). We then computed the correlation between the neural patterns for the first and second presentation of items. Half of the repeated items were identical to the first presentations (old trials), and the other half – our critical condition – were highly similar but not identical to their prior presentation (similar trials). The correlation coefficients of these pairs (r values, Pearson) were computed at the trial level and then averaged across SS and OO conditions. We hypothesized that pattern separation would require the neural patterns to become more distinct and therefore would decrease the pattern similarity in the SS condition compared to OO. We did not observe this pattern in our data (Figure 4). However, we did see a trend of condition*group interaction (F(1,81) = 2.9, p = 0.09, Figure 4-B) in CA3, highlighting the difference between the patients and controls. Consistent with our prediction, this trending interaction indicates that in patients, unlike controls, the pattern similarity in SS is marginally higher than OO (p = 0.1) suggesting an impaired pattern separation in patients. In an exploratory analysis, we found that the SS pattern similarity in patients was positively correlated (r=0.4, p = 0.018) with their “excessive pattern completion index” (see behavioral results). This finding suggests that the observed behavior of excessive pattern completion might be associated with more overlapping neural representations in CA3 leading to higher pattern similarity in SS compared to OO in patients.

**Figure 4:**
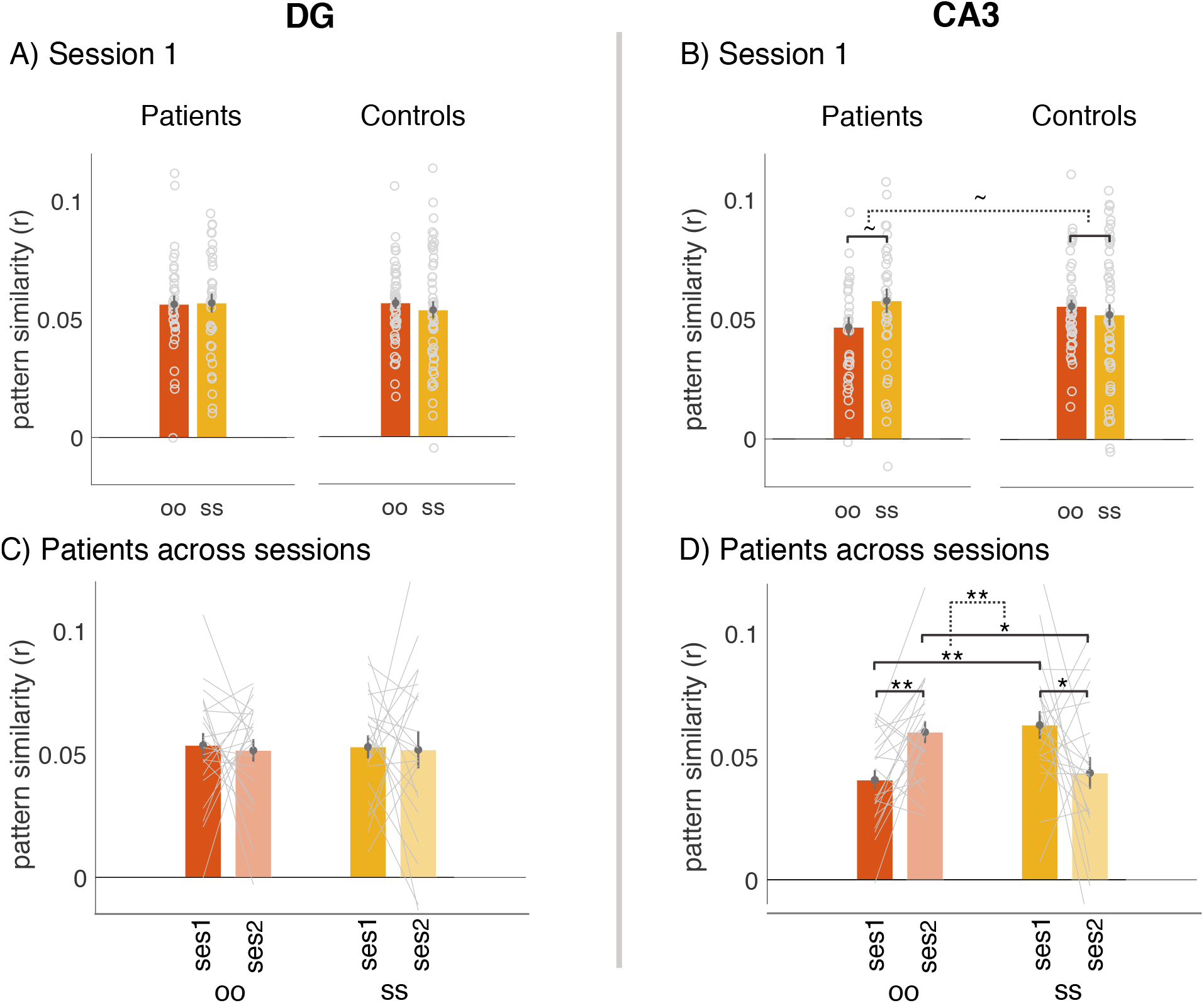
Representational similarity between the 1^st^ and 2^nd^ presentation of old and similar items during correct responses (OO, SS). **A)** Representational similarity in DG in Session 1 across the two groups **B)** Representational similarity in CA3 in Session 1 across the two groups. The dotted line depicts the trending interaction between the two comparisons. **C)** Representational similarity in DG across the two sessions in patients. **D)** Representational similarity in CA3 across the two sessions in patients. The dotted line depicts the interaction between the two comparisons.

Examining the pattern similarity in both Session 1 and Session 2 in patients who came back for the follow-up session, we found clear evidence for impaired pattern separation in CA3 in Session 1 which was reversed with treatment (Session 2). We found a significant interaction between condition and session in patients in CA3 (F(1,20) = 14.3, p = 0.001, Figure 4-D). Specifically, neural pattern similarity during SS condition was significantly higher than OO in Session 1 (p=0.002). This was reversed in Session 2, where pattern similarity during SS became significantly lower than OO (p=0.04) and resulted in the significant interaction mentioned earlier (Figure 4- D). Pairwise comparison of neural pattern similarity during SS and OO before and after treatment in CA3 revealed an increase in pattern similarity during OO (p = 0.002) and a decrease in pattern similarity during SS (p = 0.02). These results suggest that treatment flipped the direction of difference between SS and OO in patients. No main effect or interaction was observed in controls across the two sessions. DG and CA1 did not show any main effect or interaction in Session 1 in either group or across Sessions 1 and 2 (DG: Figure 4-A/C, CA1: Figure S4-B/D).

#### Trial-level analysis of neural data in hippocampal subfields

Thus far we have reported behavioral data that show reductions in both overall recognition performance as well as in fine-grained pattern separation performance in patients compared to controls. Our neural univariate data show that both DG and CA3 are differentially engaged when viewing old and similar trials in controls but not in patients, implying a relationship between activation in these regions and performance on the task. To more directly test the relationship between BOLD activation in DG and CA23 and performance, we leveraged all trials – correct and incorrect responses – and asked whether trial-by-trial variability in BOLD responses in hippocampal subfields is related to trial-by-trial behavioral responses. We specifically focused on DG during similar trials, as it has been associated with pattern separation ^27,28^, and CA3 during old trials since it has been linked to pattern completion, or retrieving a prior pattern based on a partial cue ^35,36^. We ran three trial-level mixed-effects linear models predicting the success in response to new items, old items, and similar items with univariate activity in each hippocampal subfield (Figure 5, see Methods for details of modeling) in patients and controls.

**Figure 5:**
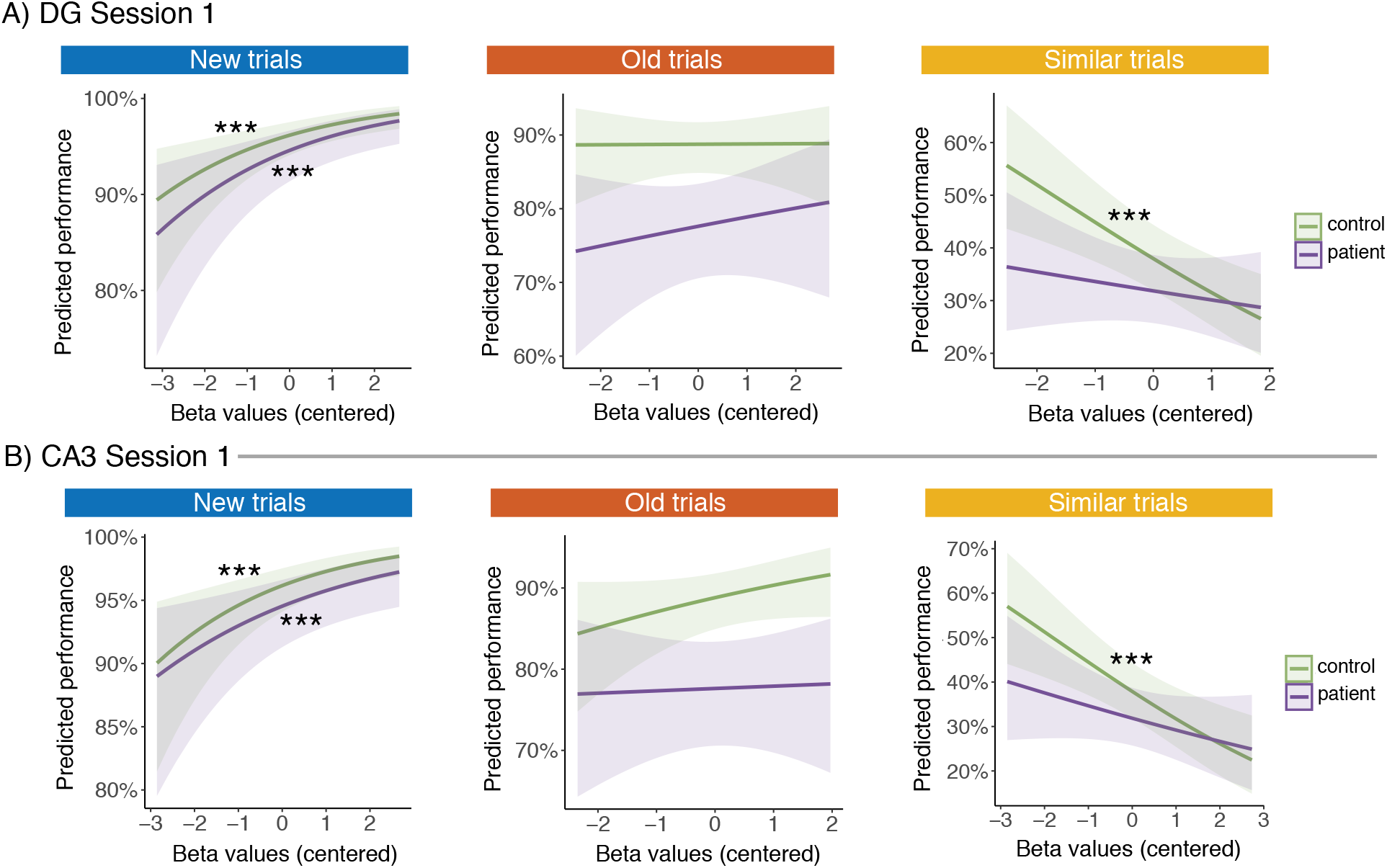
Results of trial-level mixed-effects linear models in Session 1 A) predicting the success in response to new items, old items, and similar items in DG in Session 1. B) predicting the success in response to new items, old items, and similar items in CA3 in Session 1

In DG, we found a strong positive association between trial-specific univariate activity (mean-centered beta values) and the probability of response success in both patients and controls for new trials (controls: b = 0.34, p = 0.0006, patients: b = 0.33, p = 0.001, Figure 5-A). For old trials, univariate activity in DG did not significantly predict behavior in either group. Interestingly, for similar trials, we observed a clear divergence between the results in patients and controls. In controls, more negative beta estimates (i.e. lower DG BOLD activity) correlated with a higher behavioral success rate (b = −0.28, p = 0.0005, Figure 5-A). However, this association was not present in patients where no relationship was seen between DG BOLD responses during similar trials and behavioral success. Note that this finding matches with and extends our univariate results for the SS condition (Figure 3) where we found more negative percent signal change in the control group in hippocampal subfields for the similar trial condition compared to the old condition.

We observed the same pattern of results in CA3 and CA1(Figure 5-B and S7-A). In CA3 and CA1, in both patients and controls, the trial level univariate activity during new trials predicted behavior (CA3-control: b=0.35, p = 0.0005, patient: b=0.26 p=0.006, CA1-control: b=0.50, p = 0.0001, patient: b=0.28 p=0.03) and the neural activity during similar trials were predictive of performance only in controls (CA3: b=−0.27, p= 0.0006, CA1:b=−0.29, p = 0.004), but not patients. The lack of a significant relationship between univariate activity during old trials in controls, which was seen in DG and CA1 as well, might be attributed to a ceiling effect due to high performance in these trials (only in controls).

We next asked whether these brain-behavior relationships changed across sessions during which patients were on antipsychotic medication. Similar to Session 1, modeling the response to similar trials in DG showed that the univariate response in controls predicted the behavior in Session 2 (b = −0.28, p = 0.0004). Interestingly, in Session 2 (after treatment), the association between DG univariate activity and behavioral performance during similar trials became significant in patients (b= −0.28, p = 0.02, Figure 6-A). In addition to DG, both CA3 and CA1 trial by trial univariate responses predicted successful memory of similar trials in Session 2 in controls (CA1: b = −0.29, p = 0.004 Figure S7-B, CA3:b = −0.27, p = 0.0006). In patients, however, there still was no meaningful association between univariate activity and the probability of correct responses on similar trials in CA3 and CA1. This finding suggests that treatment with antipsychotics normalized DG activation during this demanding mnemonic pattern separation task.

**Figure 6:**
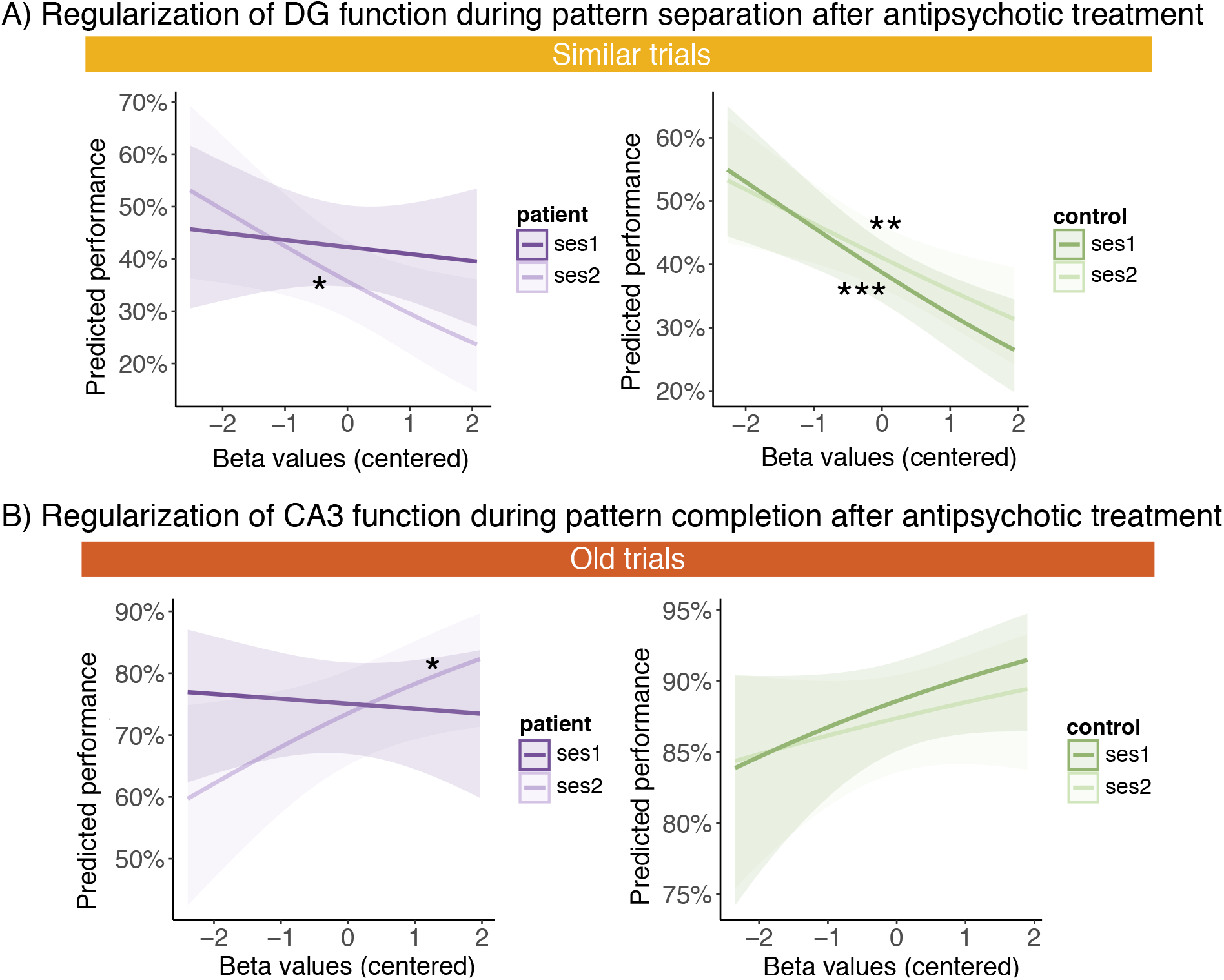
Regularization of DG and CA3 functions after antipsychotic treatment. A) predicting the success in response to *similar* items in Session 1 and Session 2 in DG. The left panel (purple) shows the model in patients. The right panel (green) presents the model results in controls. B) predicting the success in response to *old* items in Session 1 and Session 2 in CA3. The left panel (purple) shows the model in patients. The right panel (green) presents the model results in controls.

We next examined whether the brain-behavior relationships in CA3 changed across sessions in patients, over the course of treatment. Similar to Session 1, we did not observe a trial-level relationship between univariate response during old trials and behavior in controls in CA3. However, we observed a significant positive association between univariate activity during old trials and the probability of successful response in patients after treatment (b= 0.26, p = 0.034 Figure 6-B). This finding suggests a stronger role for CA3 in pattern completion mechanisms after treatment, which matches two sets of findings we previously presented in this work.

Behaviorally, we showed an excessive pattern completion in Session 1 and an improvement of this index in Session 2. In our exploratory analysis, this excessive pattern completion was higher in patients with higher SS pattern similarity. We also presented a significant interaction in multivariate responses across 2 sessions in CA3, driven by a decrease in pattern similarity in SS and an increase in pattern similarity in OO across sessions. Overall, our results suggest that treatment with antipsychotics is associated with an improvement in pattern completion mechanisms (decrease in excessive pattern completion) and is associated with changes in CA3 function measured by BOLD signal.

### Extra-hippocampal regions

Our results demonstrate aberrant hippocampal subfield recruitment (in particular DG and CA3) in patients compared to controls during trials that task mnemonic pattern separation and recognition memory in Session 1. However, behaviorally, even though patients’ pattern separation performance is impaired relative to controls, they are still able to perform the task to some extent. This raises the question that given the observed dysfunction of hippocampus, how do patients manage to perform this task? While hippocampus (and in particular area DG) has been at the center of attention in prior work on pattern separation ^41,45,53,54^, it has also been shown that cortical regions may also contribute to pattern separation ^55,56^. Therefore, the functional contribution of cortical regions to pattern separation may be relatively intact in patients, which may compensate to support successful pattern separation.

To test this hypothesis, we ran the trial-level linear mixed-effects model we previously used in hippocampal subregions (Figure 5) to model the brain-behavior relationship in “similar” trials in cortical regions (Figure 7). We selected a set of cortical regions, from early (V1) to higher level object perception and memory regions (lateral occipital and perirhinal cortex), spanning to midline default mode regions (medial prefrontal cortex and retrosplenial cortex) and executive control areas (dorsolateral prefrontal cortex). The results indicate that, in patients, response success can be predicted from univariate activity in V1 (b = 0.20, p = 9.57e−05), LOC (b = 0.19, p = 3.90e−08), perirhinal (b = 0.28, p = 0.017), mPFC (b = −0.18, p = 0.002) and dlPFC (b = 0.13, p = 0.02). Interestingly, the effects in V1 and mPFC were only observed in patients and not in controls. Retrosplenial cortex showed no effect in either group. These findings provide evidence for a compensatory role of cortical regions in pattern separation in patients.

**Figure 7:**
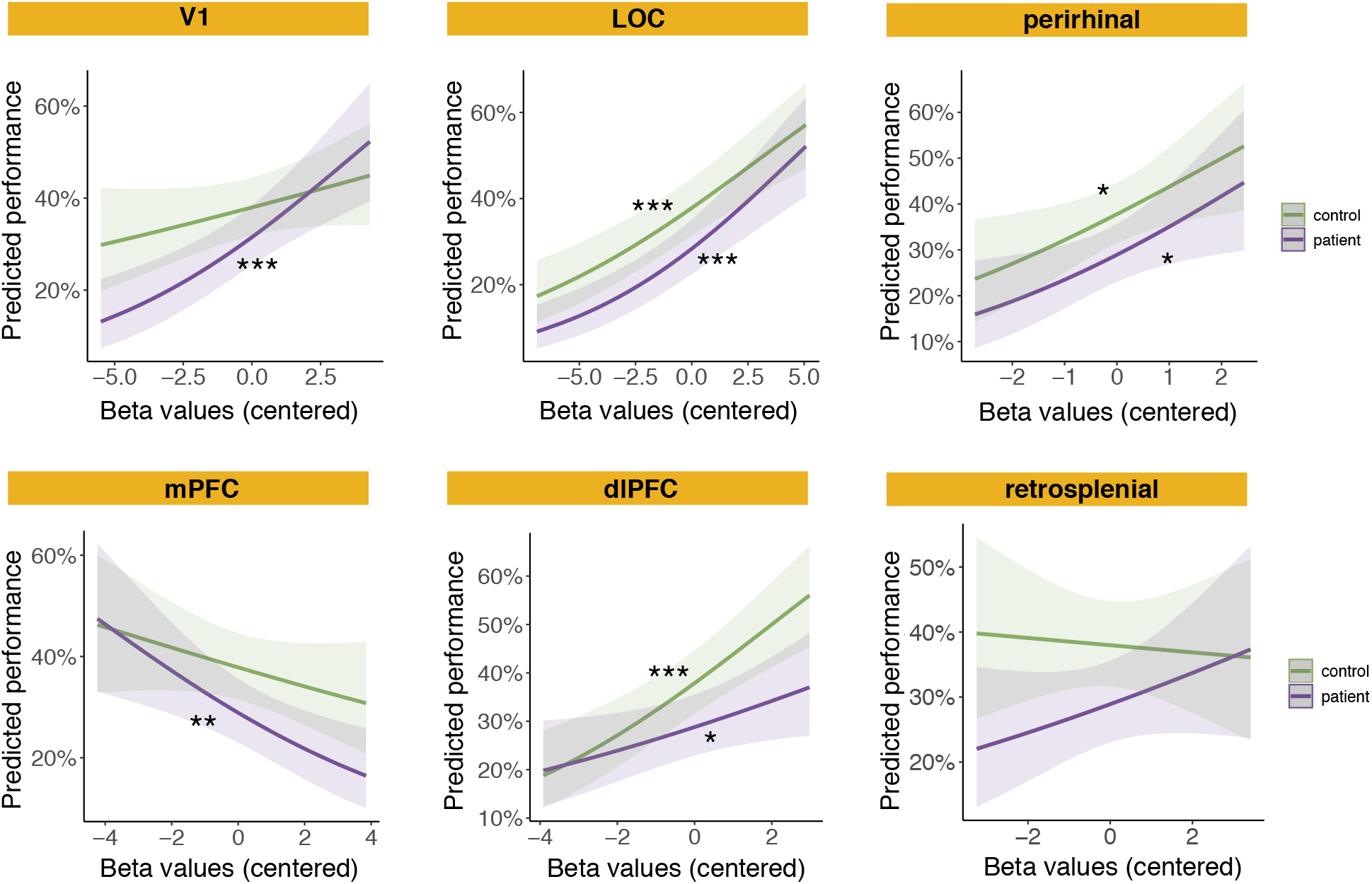
Results of trial-level mixed-effects linear models predicting the success in response to *similar* items in cortical ROIs in Session 1 (V1: primary visual cortex, LOC: lateral occipital cortex, mPFC: medial prefrontal cortex, dlPFC: dorsolateral prefrontal cortex, and retrosplenial and perirhinal cortices)

## Discussion

In the current study, using high-resolution fMRI, we examined the recruitment of hippocampal subregions and cortical regions in patients with first-episode schizophrenia performing a difficult mnemonic discrimination task before and after antipsychotic treatment. We found that patients were impaired in overall (old/similar) recognition memory as well as in their pattern separation performance and the severity of delusions in patients was positively correlated with these behavioral impairments. Analysis of univariate BOLD activity in hippocampal subfields during the task as well as its relationship with performance on a trial-by-trial basis revealed significant differences between patients and controls during the identification of similar and old items. A significant difference between BOLD activity during the successful identification of old and similar trials was observed in all hippocampal subfields in controls, whereas there was no difference in hippocampal activation in patients between these conditions. In addition, unlike controls, trial-level univariate responses during similar trials did not predict behavior in patients in any hippocampal subfield prior to treatment; the relationship between trial-level BOLD activity and performance during similar trials in DG was significant only after antipsychotic treatment. During pattern completion (recognition of old trials), CA3 trial-by-trial BOLD activation was significantly associated with memory performance after treatment in patients. Using multivariate pattern analyses, during Session 1 (pre-treatment), we found marginally greater similarity between the neural patterns of similar trials compared to old trials in patients in CA3, but this was not the case for healthy controls. Treatment inverted this pattern, resulting in significantly lower representational similarity during correct identification of similar trials than old trials. Lastly, in contrast to the impairment in hippocampal circuitry in patients during this task, trial-level BOLD response was significantly associated with performance in cortical regions during similar trials in both groups.

The main evidence that DG plays a role in pattern separation was derived from our mixed-model analyses of BOLD response before and after treatment during similar trials (mnemonic discrimination) in patients. We showed that trial-level BOLD response in hippocampal subfields could predict behavioral responses in controls both in the baseline and follow-up scan. In patients, however, there was no relationship between BOLD responses and performance in Session 1 in any of the hippocampal subfields. This relationship became stronger in all subfields after treatment and DG exhibited a significant relationship between trial-level BOLD responses and performance in Session 2. Our results suggest that treatment with antipsychotics might attenuate some DG abnormalities and extend prior work that reported treatment with antipsychotics improved hippocampal integrity in patients with schizophrenia ^40,57^.

Our main evidence for a special role of CA3 in pattern completion was obtained using trial-level mixed modeling to predict behavioral performance from trial-by-trial BOLD response during *old* trials (memory recognition). In controls, the performance was at the ceiling and thus did not provide reliable BOLD-behavior modeling results. In patients, performance was strongly impaired and was not associated with trial-level BOLD response in any of the subfields in Session 1. Only in CA3, however, performance on old trials was positively associated with BOLD responses after treatment. Furthermore, our representational similarity analysis revealed marginally greater similarity between the patterns of neural response in SS than OO in patients in CA3, but not in controls. Importantly, this increased similarity was reversed in CA3 after treatment, where the patterns of neural response in SS became less similar compared to OO, and a significant interaction between the SS and OO responses was observed across sessions. This finding supports the theory that the balance between pattern separation and pattern completion is dysregulated in individuals with schizophrenia ^1^. Moreover, it has been shown that reduced GluN1 in mouse dentate causes CA3 hyperactivity and is linked with psychosis-like behavior ^58^. Our finding of a positive correlation between neural pattern similarity during SS in CA3 and the behavioral “excessive pattern completion index” supports a possible role for CA3 hyperactivity in schizophrenia-related memory dysfunction.

Our work extended prior reports on pattern separation in healthy individuals. Prior work in healthy participants using this task has examined neural responses in a single region of interest that contained both the DG and CA3 regions, due to insufficient imaging resolution ^45^ (but see ^54^, which focuses on DG activity during pattern separation in a different task). However, we study the function of DG and CA3 as two separate regions. We found a consistent difference between the BOLD response during correct identification of similar and old trials in controls in all subfields. The lack of specificity for DG or DG/CA3 (unlike some prior work ^45,54^), suggests a synchronized response of hippocampal subfields during pattern separation, consistent with the highly connected structure of the hippocampal complex ^59^. Moreover, some earlier work on pattern separation using MST ^45^ reported a response during similar trials in the combined DG/CA3 ROI that resembled new trials (both higher than the response during repeated trials). We found a different pattern of response in controls; BOLD activity in response to similar and old trials significantly differed but similar trials also differed from new trials. This might be attributed to differences in the task design (incidental encoding vs. continuous recognition paradigm) as a prior study using the mnemonic judgment version of the task also reported lower (but not significant) response during correct identification of similar trials compared to old trials^45^. Moreover, our novel approach to analyzing the MST data using trial-by-trial modeling of the association between BOLD activity and behavior allowed us to confirm that BOLD activity during similar trials was predictive of behavior, further validating our results.

Our behavioral results are consistent with prior findings reporting a pattern separation deficit in schizophrenia ^11,12,42^ and reconcile inconsistent reports of recognition memory deficit ^11,12^. We found a large deficit in recognition memory in patients for old trials and for overall recognition of either similar or old trials. Among our behavioral measures and indices, only the “excessive pattern completion index” improved after treatment in patients and became comparable to controls. This index was calculated by summing the proportion of “old” and “similar” responses to new trials. Prior work using MST treats these two measures as general response biases. We found that the sum of these two measures was significantly higher in patients in Session 1 and improved after treatment. Moreover, patients with a higher “excessive pattern completion index” exhibited increased representational similarity during SS trials in CA3. Overall, our results indicate that NS (“similar” response to new trials) and NO (“old” response to new trials) conditions may be beyond simple general response bias and be linked to impaired pattern completion.

This is the first study reporting that pattern similarity and recognition memory performance during MST correlate with delusion severity. Specifically, we found significant relationships between pattern separation and overall recognition scores with delusion severity while participants were presented with *similar* trials -- the most taxing condition – which suggests a link between memory dysfunction and delusions. Compared to prior work on mnemonic discrimination in schizophrenia ^11,12,42^, we had a much larger patient cohort. In addition, previous work utilized BPRS ^60^ or PANSS ^61^ to quantify the positive symptoms of schizophrenia, whereas we implemented the PSYRAT scale ^62^. Furthermore, we specifically targeted delusions among the symptoms of schizophrenia, as we hypothesized that delusions may be specifically linked to memory system dysfunction. Our analysis revealed a relationship between our behavioral memory measures and PSYRAT delusions score, but not the hallucination score, consistent with our hypothesis. These findings support a link between memory dysfunction and delusions in schizophrenia and provide evidence for theories that suggest delusions are particularly linked to aberrant memories ^39^ and that memory system dysfunction contributes to psychosis ^3,14^.

Importantly, our patient cohort consisted of drug naïve or minimally treated patients newly diagnosed with schizophrenia which enabled us to study the hippocampal circuitry before it was affected by progression of the illness or prolonged antipsychotic medication. Prior work has shown more hippocampal atrophy in patients with a longer duration of psychosis ^13^. In addition, previous studies suggest that antipsychotic treatment affects the hippocampal complex ^43^. Our study provides insight into the hippocampal system in schizophrenia at the onset of the disorder, which has implications for future work on the pathophysiology of the illness.

Lastly, we found impairment in both BOLD response and representational similarity of neural patterns during *correct* identification of similar trials, when patients were able to perform the pattern separation despite a clear hippocampal deficit. We found that trial-level BOLD response during similar trials in our set of cortical ROIs (mPFC, dlPFC, V1, LOC, and perirhinal cortex) strongly predicted behavioral outcomes. Prior studies have shown intact or increased recruitment of cortical regions but impaired hippocampal activation during conscious recollection of words^21^ and associative memory ^63^ in patients with schizophrenia. Our results extend these findings to mnemonic discrimination while providing a direct association between cortical univariate activity and memory performance in response to similar trials. Considering the clear impairment in hippocampal univariate activity during correct trials (SS), our results suggest a compensatory role for cortical regions during mnemonic discrimination in patients. Furthermore, our results underscore the importance of examining not only the hippocampus but also interconnected cortical regions during mnemonic discrimination tasks ^55^.

To our knowledge, this is the first study evaluating the function of hippocampal circuitry (subfields) during a memory task in patients with schizophrenia. Through a comprehensive set of behavioral and neural analyses, the current study provides novel evidence on the impairment of the memory system in new-onset schizophrenia and the role of antipsychotic treatment

## Methods

### Participants

In this two-session fMRI study, 50 individuals newly diagnosed with schizophrenia with minimal antipsychotic exposure (mean age=23 years, range 15 −36 years, 26 females, 24 males) and 50 age-matched controls (mean age= 24 years, range18 −35 years, 28 females, 22 males) (Table 1) participated in the experiment. Data was collected at Shanghai Mental Health Center and Suzhou Guangii Hospital over 24 months. Following enrollment and screening, the first session of the experiment took place the same day or shortly thereafter (less than 10 days later). The follow-up session was performed 2-18 weeks after the first session (mean = 6.5 weeks, SD = 3.8 weeks).

**Table 1:** Demographic characteristics of participants.

|  | Healthy control<br>group<br>N = 49 | Individuals with<br>schizophrenia<br>N = 45 |  |
| --- | --- | --- | --- |
|  | Mean (*SD*) or No. | Mean (*SD*) or No. | p-value |
| Age, years | 24.6 (5.4) | 24 (6.3) | > 0.05 |
| Gender | 21 F, 24 M | 28 F, 21 M | > 0.05 |
| Handedness | 0 left | 0 left | > 0.05 |
| Education, years | 15 (2.1) | 12.6 (2.7) | < 0.0001 |
| Race / ethnicity | 49 Asian/Chinese | 45 Asian/Chinese |  |
| PSYRAT delusion score |  | 16.8 (4.1) |  |

Patients included in the study had experienced a first episode of non-affective psychosis defined by persistence of delusions for at least 4 days per week for at least 4 weeks in the absence of psychotomimetic substance use or other potential organic etiologies (including epilepsy or a history of significant head trauma). During screening and the day of imaging, they scored at least a 2 on the “Amount of preoccupation with delusions” and “Conviction” items of the Psychotic Symptom Rating Scale (PSYRATS). Patients had no prior diagnosis of major mood disorder or other Axis I disorder other than Schizophrenia, Schizoaffective Disorder, or Schizophreniform Disorder, no significant medical or neurological illness by history or physical exam including seizure disorder, history of loss of consciousness related to head trauma or developmental disorder including mental retardation and were eligible to undergo fMRI scanning. Patients included in the study had minimal or no antipsychotic treatment at the time of the experiment. At the time of the fMRI scan, 20 of 50 patients had no prior antipsychotic treatment (drug-naive group). The other 30 patients (minimally-treated group) had less than 14 days of antipsychotic treatment (mean = 3.8 days, range = 1 - 13 days). The age-matched healthy controls were free of any Axis I disorder and any history of psychosis, any psychotropic medication use in the past six months, any substance abuse in the past six months, and any medical or neurological problem that might affect hippocampal function and did not endorse delusions on the Peter’s Delusion Index.

One healthy control and four patients were excluded due to chance-level performance, and one patient was unable to complete the task. Therefore baseline behavioral analyses were performed on 45 patients and 49 controls (Session 1). Thirty-three patients and 47 healthy controls returned for the follow-up Session 2. Four patients were excluded from imaging analyses due to excessive head motion. Further, in both patient and control groups, runs with excessive head motion (framewise displacement > 0.9 in more than 10% of Repetition Times) were removed. In the univariate and multivariate analyses, only participants that had more than 7 correct trials were included in the analyses.

PhD-level psychologists or psychiatrists at Shanghai Mental Health Center and Suzhou Guangii Hospital evaluated all participants for their ability to provide consent. The study was approved by IRBs at NYU Grossman School of Medicine, Suzhou Guangii Hospital, and Shanghai Mental Health Center. Written informed consent was obtained from all participants. For participants under 18 years old the consent was obtained from their guardian.

### Study design

Participants completed a version of the Mnemonic Similarity Task^48^ across 4 runs. In this continuous recognition task, a series of images were presented on the screen and participants responded whether each image was new (“new” response), repeated (“old” response), or similar to a previous item but slightly different (“similar” response). The images were color photographs of objects ^37^. During each presentation, an image remained on the screen for 2.5 seconds against a gray background after which a blue frame appeared around the image, indicating that the response window was about to close, and the image remained on the screen for another 1 second. Participants were asked to respond as quickly as possible from the onset of the stimulus presentation. Response options (“new”, “old”, or “similar”) appeared under the image in each trial in text format. Intertrial intervals (ITIs) were jittered between 1.5, 4, and 6.5 seconds, with a gray background displaying a fixation cross. During each run, 48 new items were presented, 20 of which were repeated (“old” trials), and another 20 were similar but no identical items (“similar” trials). A total of 192 new items, 80 old items, and 80 similar items were presented over four runs. A 7-minute rest scan followed each task session, in which participants were asked to relax. In total, participants completed 4 runs of 88 trials each. In Sessions 1 and 2, the task was identical, but the stimuli set was different.

### MRI acquisition

Imaging data were obtained using a 3T Siemens Skyra scanner with a 32-channel head coil at Suzhou Guangji Hospital imaging center. Anatomical images were collected using T1 and T2 weighted protocols (1mm^3^ resolution). Both T1 and T2 images were obtained from 57 participants (28 patients and 29 controls) and only T1 images from 37 participants (17 patients and 20 controls), due to technical issues that delayed the acquisition of T2 images at the beginning of the data collection period. Functional images were acquired using a multiband EPI sequence (TR=2500ms, TE=28 ms, FOV=204mm*204mm, flip angle 75 degrees, slice thickness=2mm, and PAT=4). Field maps were collected using a double-echo gradient echo field map sequence to correct B0 inhomogeneities.

### Preprocessing

Results included in this manuscript come from preprocessing performed using *fMRIPrep* 20.2.3 ^64^ (RRID:SCR_016216), which is based on *Nipype* 1.6.1^65^ (RRID:SCR_002502).

#### Anatomical data preprocessing

A total of 2 T1-weighted (T1w) images were found within the input BIDS dataset. All of them were corrected for intensity non-uniformity (INU) with N4BiasFieldCorrection^66^, distributed with ANTs 2.3.3^67^ (RRID:SCR_004757). The T1w-reference was then skull-stripped with a *Nipype* implementation of the antsBrainExtraction.sh workflow (from ANTs), using OASIS30ANTs as target template. Brain tissue segmentation of cerebrospinal fluid (CSF), white-matter (WM) and gray-matter (GM) was performed on the brain-extracted T1w using fast (FSL 5.0.9, RRID:SCR_002823, ^68^). A T1w-reference map was computed after registration of 2 T1w images (after INU-correction) using mri_robust_template ^69^. Brain surfaces were reconstructed using recon-all (FreeSurfer 6.0.1, RRID:SCR_001847^70^), and the brain mask estimated previously was refined with a custom variation of the method to reconcile ANTs-derived and FreeSurfer-derived segmentations of the cortical gray-matter of Mindboggle (RRID:SCR_002438 ^71^).

#### Functional data preprocessing

For each of the 4 BOLD runs per subject (across all tasks and sessions), the following preprocessing was performed. First, a reference volume and its skull-stripped version were generated using a custom methodology of *fMRIPrep*. Fieldmap-based susceptibility distortion correction was performed using phase-difference maps. The BOLD reference was then co-registered to the T1w reference using bbregister (FreeSurfer) which implements boundary-based registration^72^. Co-registration was configured with six degrees of freedom. Head-motion parameters with respect to the BOLD reference (transformation matrices, and six corresponding rotation and translation parameters) are estimated before any spatiotemporal filtering using mcflirt (FSL 5.0.9 ^73^). The BOLD time-series were resampled onto the following surfaces (FreeSurfer reconstruction nomenclature): *fsnative*, *fsaverage6*. The BOLD time-series (including slice-timing correction when applied) were resampled onto their original, native space by applying the transforms to correct for head-motion. These resampled BOLD time-series will be referred to as *preprocessed BOLD in original space*, or just *preprocessed BOLD*. The BOLD time-series were resampled into several standard spaces, correspondingly generating the following *spatially-normalized, preprocessed BOLD runs*: MNI152NLin2009cAsym, MNI152NLin6Asym. First, a reference volume and its skull-stripped version were generated using a custom methodology of *fMRIPrep*. Several confounding time-series were calculated based on the *preprocessed BOLD*: framewise displacement (FD), DVARS and three region-wise global signals. FD was computed using two formulations following Power ^74^. FD and DVARS are calculated for each functional run, both using their implementations in *Nipype* (following the definitions by Power^71^). The three global signals are extracted within the CSF, the WM, and the whole-brain masks. Additionally, a set of physiological regressors were extracted to allow for component-based noise correction (*CompCor*, ^75^). Principal components are estimated after high-pass filtering the *preprocessed BOLD* time-series (using a discrete cosine filter with 128s cut-off) for the two *CompCor* variants: temporal (tCompCor) and anatomical (aCompCor). tCompCor components are then calculated from the top 2% variable voxels within the brain mask. For aCompCor, three probabilistic masks (CSF, WM and combined CSF+WM) were generated in anatomical space. The implementation differs from that of Behzadi et al. in that instead of eroding the masks by 2 pixels on BOLD space. This mask is obtained by dilating a GM mask extracted from the FreeSurfer’s *aseg* segmentation, and it ensures components are not extracted from voxels containing a minimal fraction of GM. Finally, these masks are resampled into BOLD space and binarized by thresholding at 0.99 (as in the original implementation). Components are also calculated separately within the WM and CSF masks. For each CompCor decomposition, the *k* components with the largest singular values are retained, such that the retained components’ time series are sufficient to explain 50 percent of variance across the nuisance mask (CSF, WM, combined, or temporal). The remaining components are dropped from consideration. The head-motion estimates calculated in the correction step were also placed within the corresponding confounds file. The confound time series derived from head motion estimates and global signals were expanded with the inclusion of temporal derivatives and quadratic terms for each^76^. Frames that exceeded a threshold of 0.5 mm FD or 1.5 standardised DVARS were annotated as motion outliers. All resamplings can be performed with *a single interpolation step* by composing all the pertinent transformations (i.e. head-motion transform matrices, susceptibility distortion correction when available, and co-registrations to anatomical and output spaces). Gridded (volumetric) resamplings were performed using antsApplyTransforms (ANTs), configured with Lanczos interpolation to minimize the smoothing effects of other kernels^77^. Non-gridded (surface) resamplings were performed using mri_vol2surf (FreeSurfer). For more details of the pipeline, see the section corresponding to workflows in *fMRIPrep*’s documentation.

### Atlas and ROI definition

Freesurfer’s automated cortical and subcortical parcellation^78^ was performed as a part of our preprocessing pipeline using fMRIprep. We extracted hippocampal, perirhinal, and lateral occipital cortex (LOC) regions of interest from Freesurfer parcellations in each participant. In each participant, dentate gyrus (DG), CA1, and CA3 ROIs were created using Freesurfer’s automated hippocampal subfield segmentation (FreeSurfer 6.0.1^70^). The CA3 ROI was created by combining CA2 and CA3 as a single ROI. The procedure was performed using T1 and T2 anatomical images (only T1 images for participants who did not have T2 images). All ROIs obtained from Freesurfer were then resampled to match the resolution of the EPI images. A winner-take-all method was used to binarize the subfield ROIs to ensure no overlap between hippocampal subfields voxels^79^. The medial prefrontal cortex (mPFC), dorsolateral prefrontal cortex (dlPFC), retrosplenial, and V1 regions of interest were extracted from an atlas of functional parcellation generated based on whole-brain functional connectivity^80^.

### Behavioral analysis

The proportion of responses to each of the 9 task-response conditions was calculated after removing the trials where participants failed to respond (the number of responses out of 192 new trails mean(SD): patients = 143(50), controls = 168(32) p = 0.004, the number of responses out of 80 old trials: patients = 66(16), controls = 76(6) p = 0.0001, the number of responses out of 80 similar trials: patients = 63(18), controls = 74(8) p = 0.0003). Two-sample t-test was used to compare the proportion of responses between patient and control groups. Paired t-test was used to compare the data across two sessions within participants.

For behavioral analysis (Figure 2), all the results presented from Session 1 were obtained from all participants included in the study (patients N = 45, controls N = 49). In the analysis across sessions, only participants who returned after 2-12 weeks and completed both sessions were included (patients N = 29, controls N = 46).

### Clinical symptoms severity measure

Psychotic Symptom Rating Scales (PSYRATS) were administered to patients upon enrollment in the study and at the follow-up session. The scale consists of 16 items, 10 of which relate to hallucinations and 6 to delusions. A delusions score was created by summing ratings on questions related to delusions (the main PSYRAT score used in our analyses). Using the hallucination-related questions, we obtained an additional hallucination score.

### fMRI analysis

#### GLM modeling of correct trials

An estimation of neural response to conditions of interest was obtained using a general linear model (GLM) with seven main regressors. One regressor modeled the new trials when the response was correct (NN), the old trials when the response was correct (OO), the similar trials when the response was correct (SS), the similar trials with “new” response (SN), the similar trials with “old” response (SO), the new trials when the response was incorrect (NS and NO), and the old trials when the response was incorrect (ON and OS). Regressors were modeled with an event duration of 3.5 seconds using a boxcar and were convolved with a double-gamma hemodynamic response function (HRF). Modeling was implemented using FSL FEAT^81^. Trials with framewise displacement > 0.9 were flagged as confound (motion outliers) in feat. Nuisance regressors in the model included six head motion parameters, their 1^st^ temporal derivatives, the top 5 anatomical CompCor estimates for white matter (WM), and the top 5 anatomical CompCor estimates for cerebrospinal fluid (CSF) (see functional preprocessing for details about extracting aCompCor estimates). Modeling was performed on spatially smoothed data (5mm FWHM).

Due to the difficulty of the task during similar trials, there were no correct responses to similar trials (SS) in some participants/runs. Those runs were modeled with empty events and were excluded from level 2 GLM. Ultimately, the subsequent univariate ROI analysis used the conditions for correct responses (NN, OO, and SS) that were modeled with 7 or more events across runs. Featquery (FSL) was performed to obtain the percent signal change in each region of interest. The main effects and interactions of the SS and OO measures across patient and control groups were calculated using a two-way mixed ANOVA. The statistical significance of the changes across the two sessions in each group was calculated using a 2×2 repeated measure ANOVA. Specific differences between conditions and groups were obtained using post-hoc pairwise comparisons (t-test) and p-values were adjusted using Bonferroni correction.

#### Least Square Separate (LSS) modeling of all trials

To obtain the estimates of brain response in each trial, a separate GLM model was conducted for each trial (LSS approach ^52,82^). Each trial was modeled using a regressor of interest (the event including that trial) and 3 regressors of no interest (all “new” trials, all “old” trials, and all “similar” trials, excluding the current event from the corresponding group). The model used boxcars of 3.5 seconds convolved with HRF. Nuisance regressors in the model included six head motion parameters, their 1^st^ temporal derivatives, the top 5 anatomical CompCor estimates for white matter (WM), and the top 5 anatomical CompCor estimates for cerebrospinal fluid (CSF). Modeling was performed on non-smoothed data. This method was used to estimate trial-level parameter estimates for multivariate analysis.

#### Multivariate analysis

Representational similarity analysis^83,84^ was performed by correlating the vector of beta values obtained from each ROI for the first and second presentation of items (Pearson). Correlation values in SS and OO conditions were averaged to obtain overall values. The statistical differences between group comparisons were obtained using a two-way mixed ANOVA, and the effects across sessions were measured by a 2×2 repeated measure AVONVA. P-values obtained from post-hoc comparisons were adjusted using Bonferroni correction.

Our univariate and multivariate analyses (Figures 3 and 4) were focused on the conditions with *correct* responses, we included only a subset of participants who had enough correct trials across the entire experiment in the SS condition (all participants included in the imaging analysis had enough OO trials, min = 7, mean = 60 trails). In each section, we first compared the patients (N = 34) and controls (N = 49) in Session 1. Then, we compared the two sessions in the subset of participants who completed both sessions in each group (patients N = 22 and controls N = 46).

#### Mixed-Effects Models

We used trial-level mixed-effects linear modeling to examine the relationship between the univariate brain response and behavior. As this analysis is not limited to correct trials, we were able to include all participants and runs with acceptable image quality (patient N = 41, control N = 49). Participants who completed both sessions were included in the analyses across sessions (patient N = 28, control N = 46). In this model, beta estimates from the LSS model were extracted in each ROI, were scaled (divided by 100), and mean-centered in each group/condition. Beta values +/- 3.5 standard deviations were removed in each subject and ROI. The resulting values were used as trial-level beta estimates in the model. Trials in which participants failed to respond were removed before modeling. Models were implemented using glmer in the R package. We ran two sets of models. One in patients and controls in Session 1 with the following formula: Memory performance ∼ beta values * group + (1 | subjects)

The second models included the data from Session 1 and Session 2 and were modeled separately in patients and controls with the following formula:

Memory performance ∼ beta values * session + (1 | subjects)

## Supplementary Material

### Supplementary results

#### Drug native vs. minimally treated

Patients in our study were newly diagnosed with schizophrenia and had received any antipsychotic medication for less than 2 weeks prior to the experiment (mean = 3.8 days, median = 3, min = 1, max = 13). While full antipsychotic effects may be delayed over many weeks, dopamine D2 receptor blockade occurs rapidly after administration and may affect both symptoms and memory acutely^85^. Therefore, before combining medication-naïve and minimally-treated patients into one group (to gain more experimental power), we ran some additional analyses to ensure that antipsychotic medication exposure prior to Session 1 was not associated with a difference in memory function in the minimally treated group compared to the medication-naïve group. We split the patient group data in Session 1 into the ‘drug naïve’ (N = 17) and ‘minimally treated’ (N = 28) subgroups (See Methods) and recalculated the recognition memory and pattern separation indices (Figure S2-A). The two groups did not differ in recognition memory index. The minimally treated group had a marginally higher pattern separation index compared to the drug naïve group (t(43) = −1.8014, p = 0.078, CI = [−0.23 0.013]). However, the number of days on treatment and the pattern separation index of the minimally treated group were not significantly correlated (r = 0.23, p = 0.23). The drug naïve and minimally treated subgroups also did not differ in their PSYRAT scores (t(43) = 1.04, p = 0.3). Furthermore, there was no association between the number of days on treatment and the PSYRAT score in the minimally treated group (r = −0.03, p = 0.86). Given these observations, we combined these two subgroups into one patient group to obtain more power in our neural analyses.

#### Memory behavior in Session 2

We compared the memory indices in Session 2 between patients and controls (Figure S3). Controls showed higher values in all indices, however, there was no significant difference between pattern separation index of the two groups (t(73) = 1.12, p = 0.26). Recognition memory (t(73) = 2.80, p = 0.006, CI = [0.04 0.23]), overall recognition of old trials (t(73) = 2.92, p = 0.004, CI = [0.03 0.18]), and overall recognition of similar trials (t(73) = 2.57, p = 0.01, CI = [0.02 0.20]) were significantly lower in patients. Additionally, we repeated these comparisons in Session 1 in this subset of participants who completed both sessions. We found a trending lower pattern separation index in patients (t(73) = 1.4, p = 0.16) in Session 1. Recognition memory (t(73) = 3.4, p = 0.0009, CI = [0.06 0.24]), overall recognition of old trials (t(73) = 2.92, p = 0.0003, CI = [0.07 0.24]), and overall recognition of similar trials (t(73) = 3.18, p = 0.002, CI = [0.05 0.23]) were significantly lower in patients. Note that this analysis has less power compared to what we previously reported for Session 1 (Figure 2), as we included only the subset of participants who completed both sessions. Comparing the indices within each group, we found no significant difference between Session 1 and Session 2 values in patients or controls (Figure S3). Pattern separation index was marginally higher in Session 2 compared to Session 1 in controls (t(45) = 1.48, p = 0.14, CI = [−0.01 0.07]).

---

**Figure S1:**
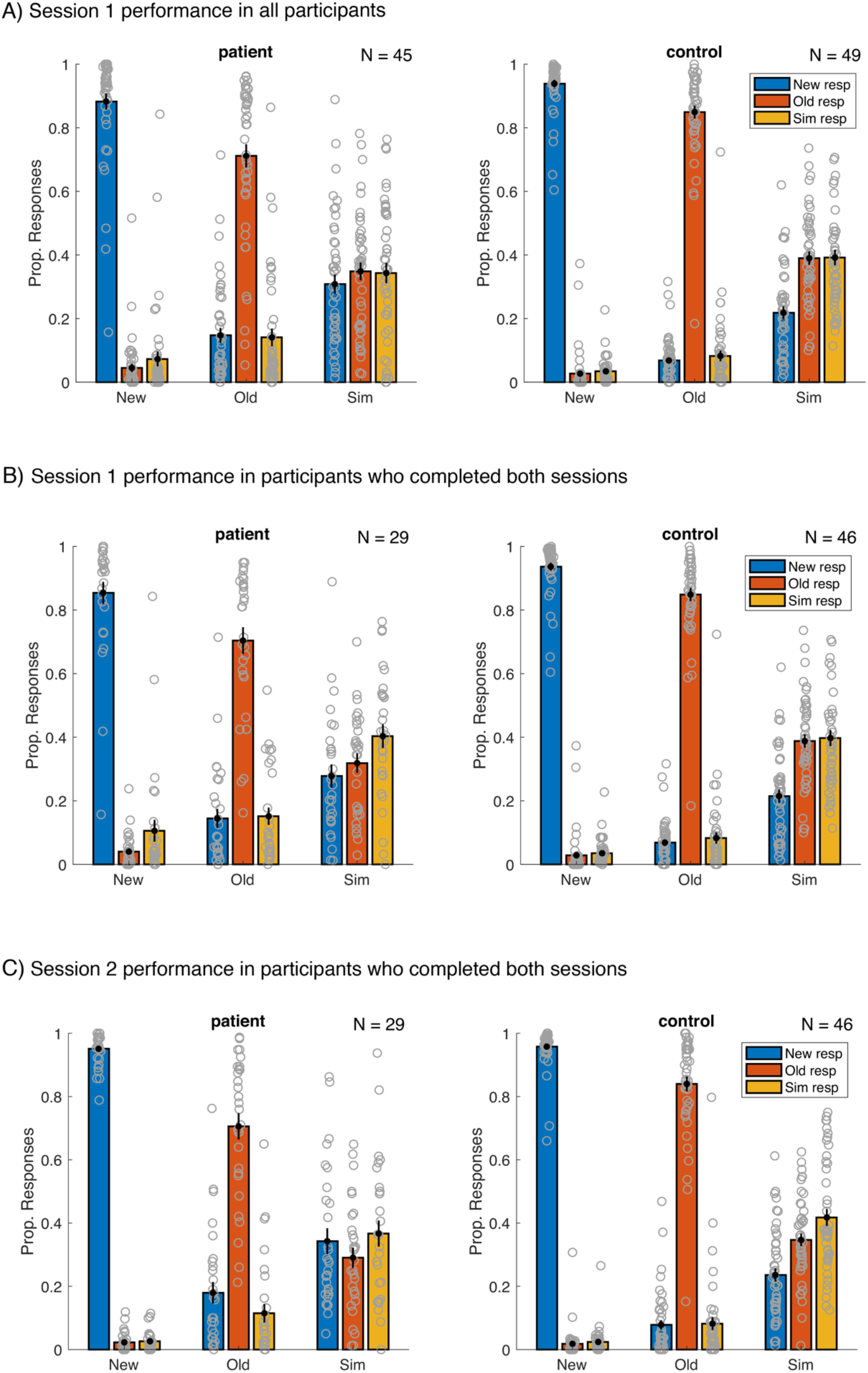
Proportion responses to each stimulus type (new, old, similar) in patients and controls. A) depicts the proportion of responses in Session 1 in all participants. B) depicts the proportion of responses in Session 1 in participants who returned after 2-18 weeks and completed the Session 2 experiment. C) depicts the proportion of responses in Session 2 in participants who returned after 2-18 weeks and completed the Session 2 experiment.

**Figure S2:**
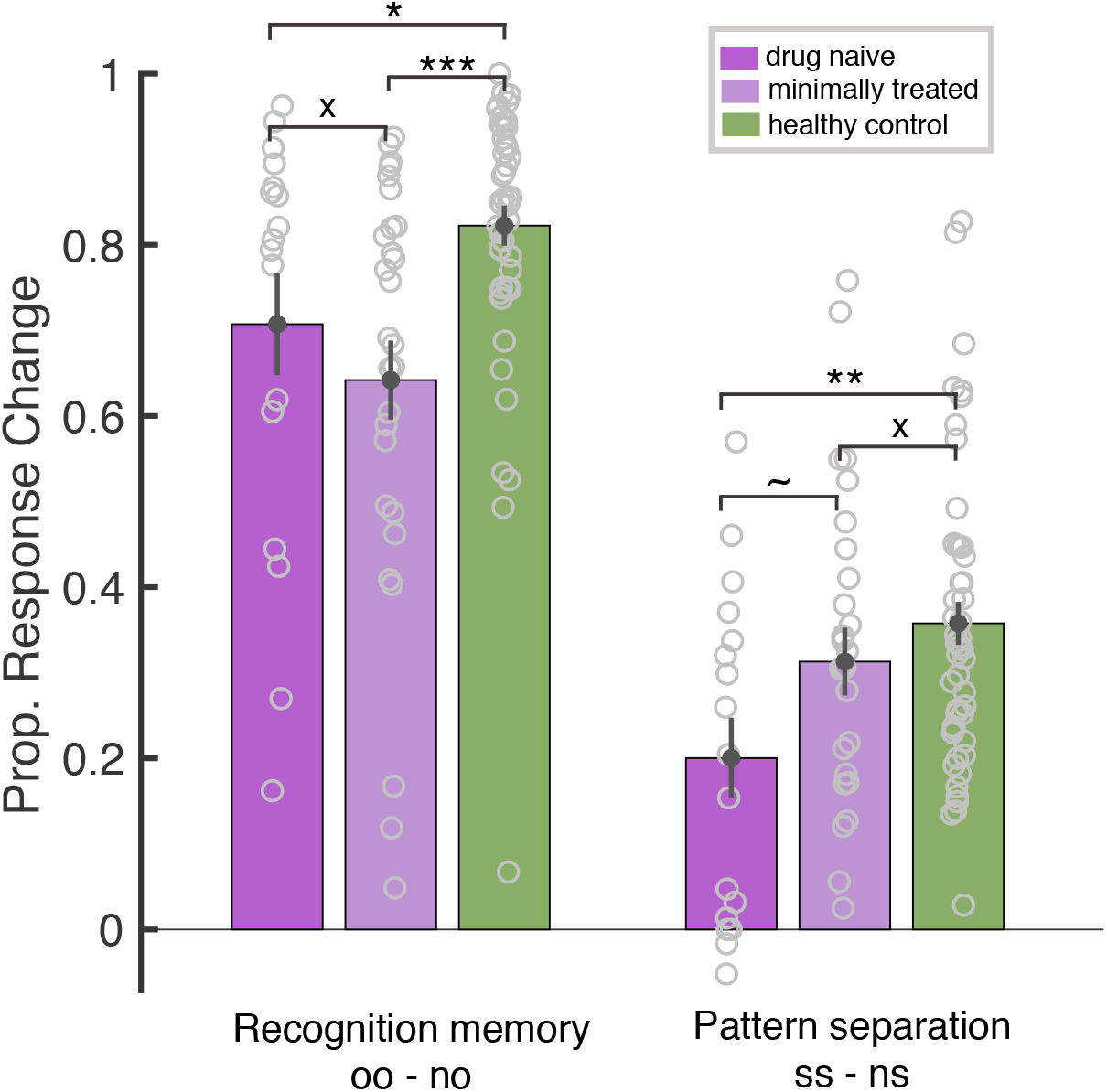
Recognition memory and pattern separation indices in Session 1 across drug subgroups and controls.

**Figure S3:**
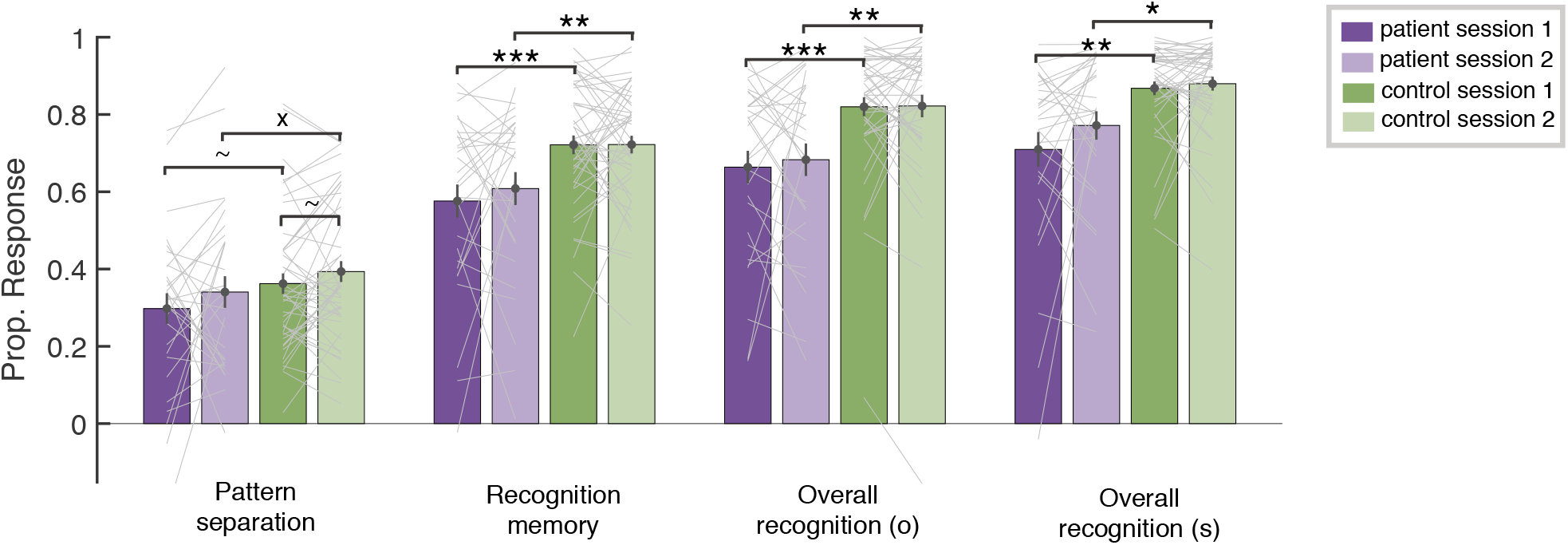
Comparing memory indices between Session 1 and Session 2 in participants who completed both sessions (patients N = 29, controls N = 46). Pattern separation, recognition memory, overall recognition of old items, and overall recognition on similar items are shown in patients and controls in Session 1 and Session 2.

**Figure S4:**
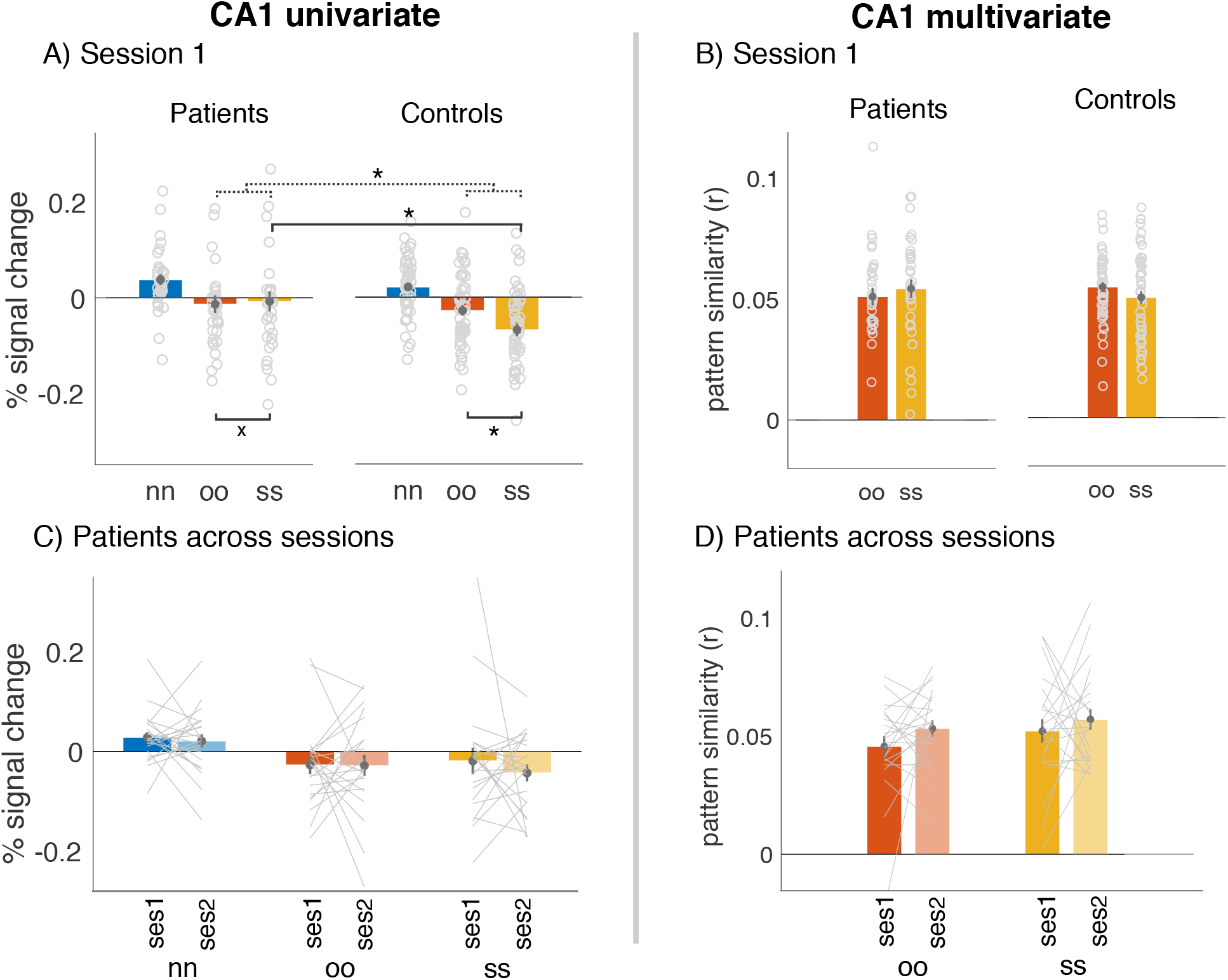
Depicts univariate and multivariate results in CA1. Dotted lines depict the interaction between the two linked comparisons. A) Percent signal change in CA1 in Session 1 across the two groups Representational similarity in CA1 in Session 1 across the two groups. C) Percent signal change in CA1 across the two sessions in patients. D) Representational similarity in CA1 across the two sessions in patients.

**Figure S5:**
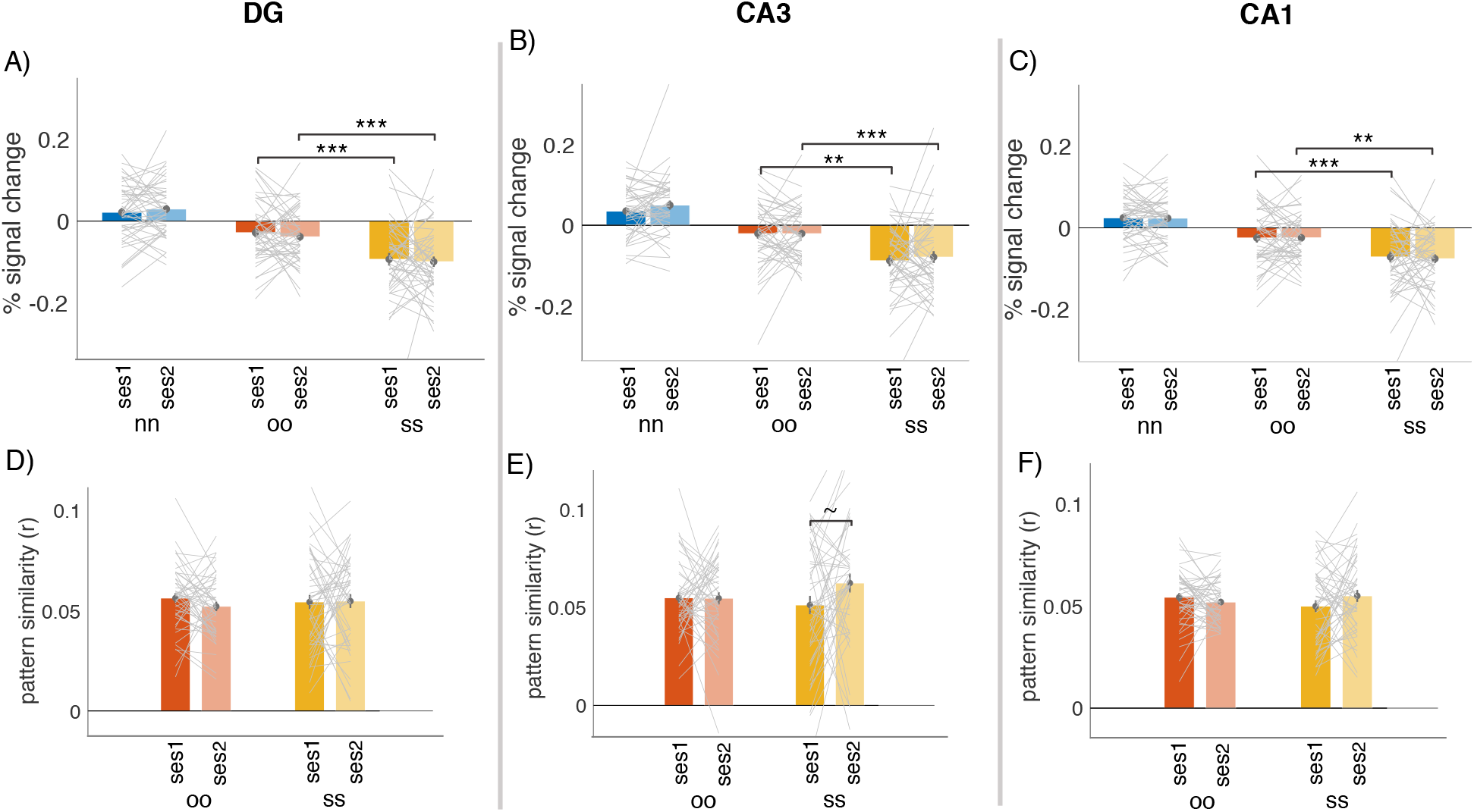
Depicts univariate and multivariate results across the two sessions in controls. A) Percent signal change in DG across the two sessions in controls. B) Percent signal change in CA23 across the two sessions in controls. C) Percent signal change in CA1 across the two sessions in controls. D) Representational similarity in DG across the two sessions in controls. E) Representational similarity in CA23 across the two sessions in controls. F) Representational similarity in CA1 across the two sessions in controls.

**Figure S6:**
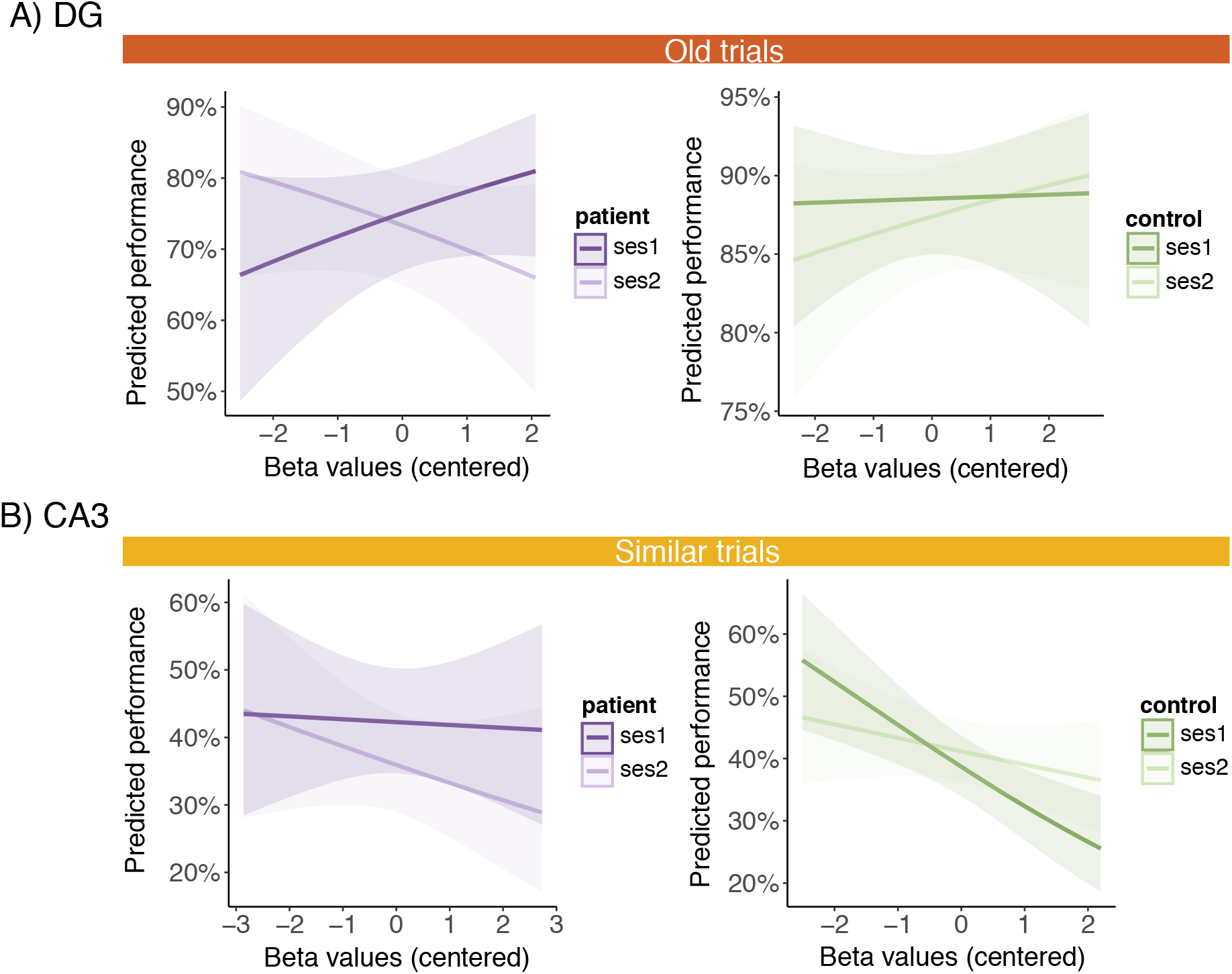
Results of trial-level mixed-effects linear models across session **A)** predicting the success in response to *old* items in Session 1 and Session 2 in DG. **B)** predicting the success in response to *similar* items in Session 1 and Session 2 in CA3.

**Figure S7:**
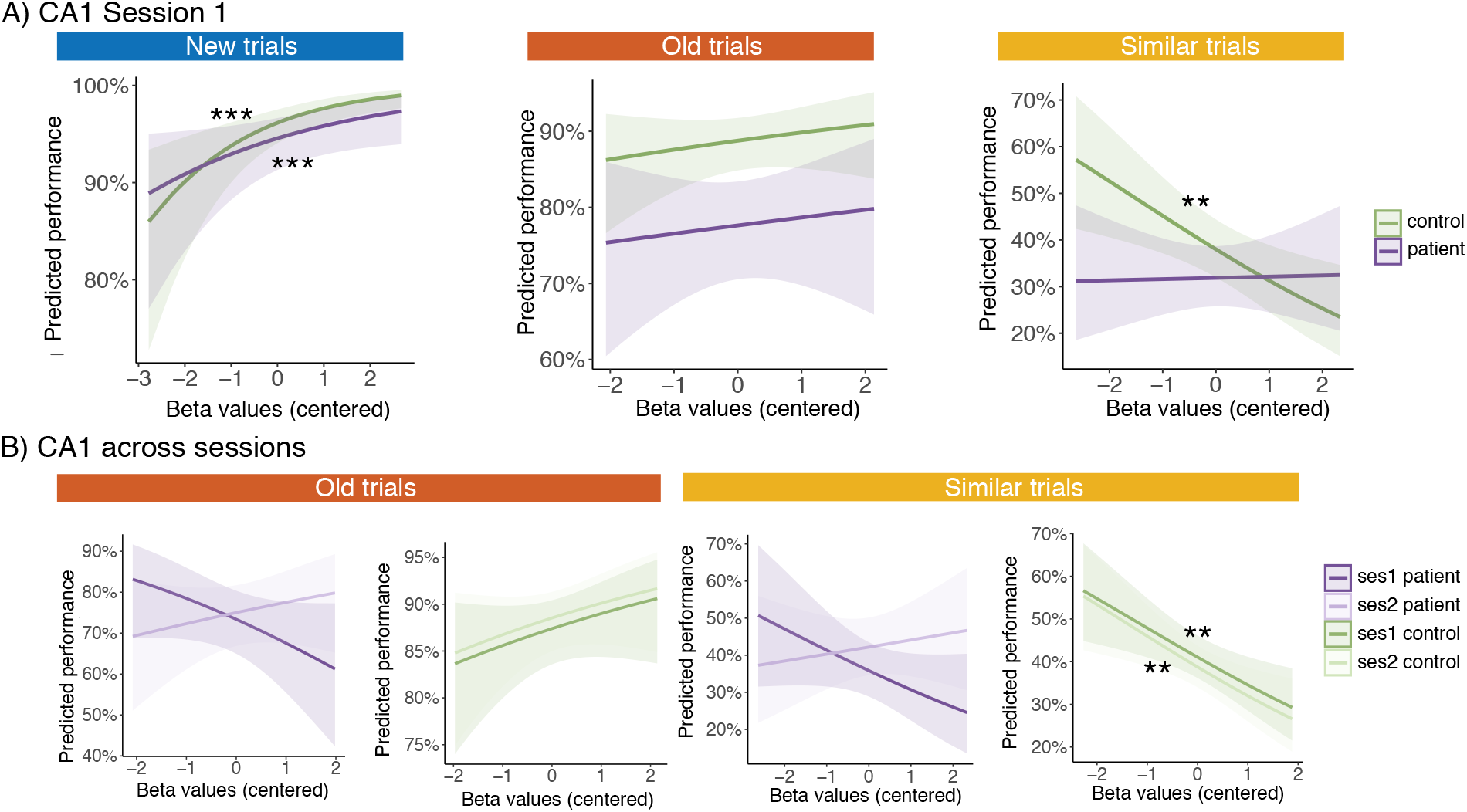
Results of trial-level mixed-effects linear models **A)** predicting the success in response to new items, old items, and similar items in CA1in Session 1 **B)** predicting the success in response to *old* and *similar* items in Session 1 and Session 2 in CA1.

